# Structural basis of transcription antitermination by Qλ: NusA induces refolding of Qλ to form nozzle for RNA polymerase exit channel

**DOI:** 10.1101/2022.03.25.485794

**Authors:** Zhou Yin, Jeremy G. Bird, Jason T. Kaelber, Bryce E. Nickels, Richard H. Ebright

**Author notes:** Corresponding Author: Richard H. Ebright.

## Abstract

Lambdoid bacteriophage Q proteins are transcription antipausing and antitermination factors that enable RNA polymerase (RNAP) to read through pause and termination sites. Q proteins load onto RNAP engaged in promoter-proximal pausing at a Q binding element (QBE) and adjacent sigma-dependent pause element to yield a Q-loading complex, and translocate with RNAP as a pausing-deficient, termination-deficient Q-loaded complex. In previous work, we showed that the Q protein from bacteriophage 21 (Q21) functions by forming a “nozzle” that narrows and extends the RNAP RNA-exit channel, preventing formation of pause and termination RNA hairpins. Here, we report atomic structures of four states on the pathway of antitermination by the Q protein from bacteriophage λ (Qλ), a Q protein that shows no sequence similarity to Q21 and that, unlike Q21, requires the transcription elongation factor NusA for efficient antipausing and antitermination. We report structures of Qλ, the Qλ-QBE complex, the NusA-free “pre-engaged” Qλ-loading complex, and the NusA-containing “engaged” Qλ-loading complex. The results show that Qλ, like Q21, forms a nozzle that narrows and extends the RNAP RNA-exit channel, preventing formation of RNA hairpins. However, the results show that Qλ has no three-dimensional structural similarity to Q21, employs a different mechanism of QBE recognition than Q21, and employs a more complex process for loading onto RNAP than Q21, involving recruitment of Qλ to form a “pre-engaged” loading complex, followed by NusA-facilitated refolding of Qλ to form an “engaged” loading complex. The results establish Qλ and Q21 are not structural homologs and are solely functional analogs.

**SIGNIFICANCE STATEMENT:** Bacteriophage Q proteins are textbook examples of regulators of gene expression that function at the level of transcription antitermination. Here, we report structures defining the mechanism of antitermination by the Q protein of bacteriophage λ (Qλ). The results show Qλ forms a “nozzle” that narrows and extends the RNA polymerase RNA-exit channel, precluding the formation of terminator RNA hairpins. The results show Qλ exhibits no structural similarity to the Q protein of bacteriophage 21 (Q21), employs a different mechanism for DNA binding than Q21, and employs a more complex process of loading onto RNA polymerase than Q21. We conclude Qλ and Q21 are not structural homologs and are solely functional analogs, akin to a bird wing and a bat wing.

## INTRODUCTION

Lambdoid bacteriophage Q proteins are transcription antitermination and antipausing factors that enable RNA polymerase (RNAP) to read through pause and termination sites (1–7), reviewed in (8–10).

Q proteins load onto transcription elongation complexes (TECs) engaged in promoter-proximal pausing to yield “Q-loading complexes,” and Q proteins subsequently translocate with TECs as pausing-deficient, termination-deficient “Q-loaded complexes” (3–10).

The Q-dependent gene regulatory cassette consists of the gene for Q, followed by a transcription unit comprising a promoter (PR′), a promoter-proximal σ-dependent pause element (SDPE), a terminator, and downstream genes [Fig. 1A; (3–10)]. In the absence of Q, RNAP initiating transcription at the PR′ promoter pauses at the SDPE and terminates at the terminator, and, as a result, fails to transcribe downstream genes. In the presence of Q, RNAP initiating at the PR′ promoter rapidly escapes the SDPE and reads through the terminator, and, as a result, transcribes downstream genes (3–10).

**Fig. 1.**
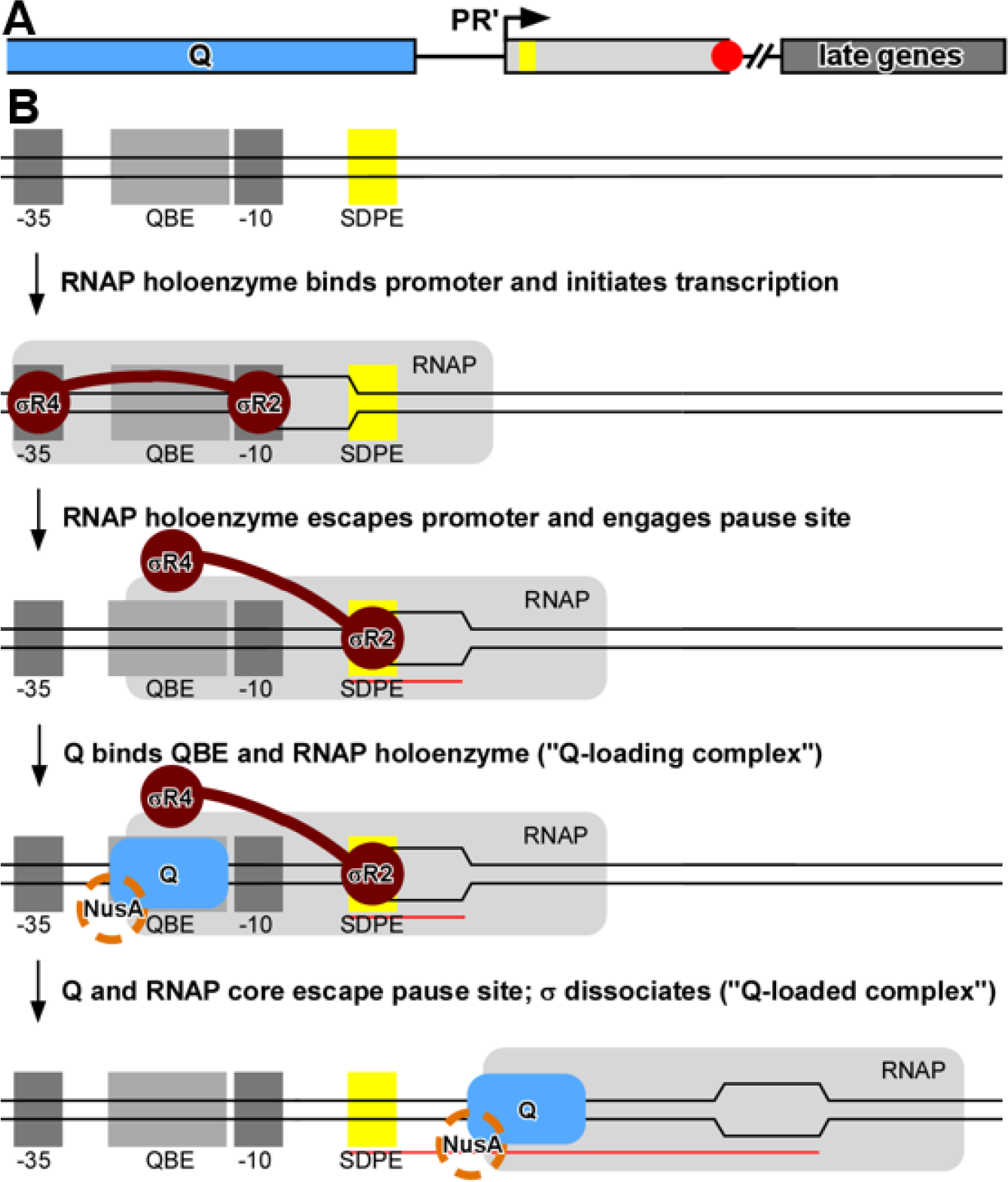
Biological function of Q. **(A)** Q-dependent regulatory cassette, consisting of gene for Q (blue) and adjacent transcription unit comprising PR’ promoter (arrow), SDPE (yellow rectangle), terminator (red octagon), and bacteriophage late genes (gray). **(B)** Steps in assembly and function of a Q-dependent transcription antitermination complex. Promoter -35 and -10 promoter elements, dark gray rectangles; QBE, light gray rectangle; SDPE, yellow rectangle; RNAP core enzyme, light gray; σ, brown; Q, blue; NusA [needed for efficient antitermination by Qλ and Q82; (2, 4, 51, 52)], orange; DNA nontemplate and template strands, black lines (unwound transcription bubble indicated by raised and lowered line segments); RNA, red line.

Q functions at the Q-dependent gene regulatory cassette by first forming a Q-loading complex, comprising a Q protein bound to a Q binding element (QBE) and a σ-containing TEC paused at the SDPE (Fig. 1B, lines 1-4), and then forming a Q-loaded complex, comprising a Q-containing TEC that processively, over thousands of base pairs, ignores pause and termination sites [Fig. 1B, line 5; (3–11)].

Q proteins comprise three protein families: the Q21 family (Pfam PF06530; 5,251 entries in NCBI), the Qλ family (Pfam PF03589, 7,904 entries in NCBI), and the Q82 family (Pfam PF06323, 2,635 entries in NCBI) (12). Q proteins from the three protein families exhibit equivalent antitermination and antipausing activities, perform equivalent regulatory functions, and are encoded by genes that exhibit equivalent positions in bacteriophage genomes (8, 12–15), but Q proteins from the three protein families exhibit no obvious sequence similarity (8, 12–15). This raises the question whether Q proteins from the three families possess three-dimensional structural homology despite the absence of obvious sequence similarity, or whether, instead, they lack three-dimensional structural homology and are solely functional analogs (akin to a bird wing, a bat wing, and a fly wing).

We recently reported a set of structures that defined the structural basis of antitermination and antipausing by the Q protein of lambdoid bacteriophage 21 (Q21): i.e., Q21, the Q21-QBE complex, the Q21-loading complex, and the Q21-loaded complex [(12), see also (16)]. In the Q21-QBE complex, two Q21 protomers interact with two tandem, directly repeated, DNA subsites (12, 16). In the Q21-loading complex, one of the two Q21 protomers that interacts with the QBE also interacts with a σ-containing TEC, forming a “Q torus,” or “Q nozzle,” that narrows and extends the RNAP RNA-exit channel (12, 16). In the Q21-loaded complex, the nascent RNA product threads through the Q nozzle, preventing the formation of pause and termination RNA hairpins, and topologically linking Q to the TEC, yielding an essentially unbreakable, processively acting, antipausing and antitermination complex (12, 16).

Here, we report a set of structures that define the structural basis of antitermination by the Q protein of bacteriophage λ (Qλ): i.e., a crystal structure of Qλ, a crystal structure of a Qλ-QBE complex, a cryo-EM structure of a “pre-engaged” Qλ-loading complex, and a cryo-EM structure of a NusA-containing, “engaged” Qλ-loading complex.

The results reveal that Qλ, like Q21, forms a nozzle that narrows and extends the RNAP RNA-exit channel. The results further reveal that the three-dimensional structures, the mechanisms of QBE recognition, and the mechanisms of Q loading differ for Qλ and Q21, and thus that Qλ and Q21 are not structural homologs and are solely functional analogs (akin to a bird wing and a bat wing).

## RESULTS

### Structure of Qλ

In previous work, we determined a crystal structure at 2.1 Å resolution of a Qλ fragment lacking part of the 60-residue intrinsically disordered N-terminal segment of Qλ [Qλ^39-207^; (17)]. In the present work, we have determined crystal structures having higher resolutions (1.46 Å for a crystal structure in the same crystal form; 1.97 Å for a crystal structure in a new crystal form) of a Qλ fragment lacking the entire 60-residue intrinsically disordered N-terminal segment and having a substitution that increases Qλ-QBE binding affinity [Qλ^61-207;N61S;E134K^; (18); Fig. 2; Table S1].

**Fig. 2.**
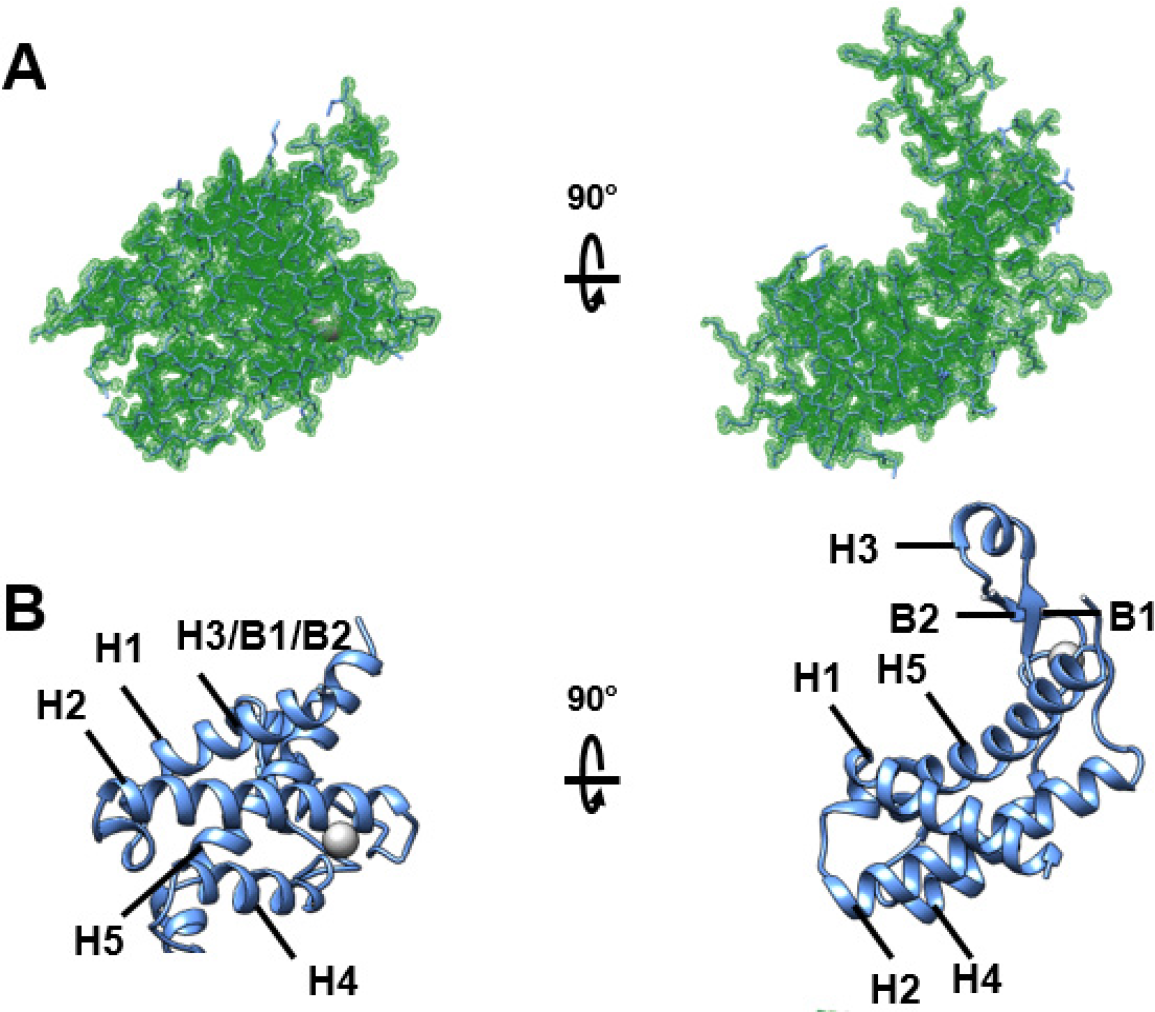
Structure of Q**λ**. **(A)** Structure of Qλ (type-I crystal structure; two orthogonal views). Experimental electron density map (green mesh, 2mFo-DFc map contoured at 1.0σ) and fitted atomic model (blue). **(B)** Structure of Qλ (type-I crystal structure; ribbon representation; two orthogonal views). Gray sphere, Zn^2+^.

The new structures confirm that Qλ comprises a globular domain containing a canonical helix-turn-helix (HTH) DNA binding motif (“Qλ body”; residues 61-113 and 153-207) and a type I shuffled (19) zinc ribbon (“Qλ arm”; residues 114-152; Fig. 2). The new structures also confirm that Qλ exhibits no three-dimensional structural similarity to Q21, apart from the presence in Qλ of a canonical HTH motif, and presence in Q21 of a non-canonical, interrupted helix-turn-loop-helix motif [HT[loop]H; (12, 16)].

### Structure of Qλ-QBE complex

We have determined a crystal structure of the Qλ-QBE complex at 2.18 Å resolution, by use of single-wavelength anomalous dispersion (Qλ^61-207;N61S;E134K^-QBE; Figs. 3, S1; Table S1). The structure shows that Qλ interacts as a monomer, in an extended conformation, with a DNA site that spans more than one turn of DNA (13 bp; Figs. 3, S1A). The Qλ body interacts, through its HTH motif, with the DNA major groove at positions 1-4 of the DNA site (Fig. 3A-B,D-E), and the Qλ arm interacts, through residues close to the residues that coordinate the Zn^2+^ ion of its zinc ribbon, with the DNA major groove at positions 11-13 of the DNA site (Fig. 3A-B,F). The observed interactions are consistent with, and account for, genetic and biochemical results defining Qλ and QBE determinants for Qλ-QBE interaction [Fig. S1A; (4, 18, 20)]. The E134K substitution that increases DNA-binding affinity (18) replaces a negatively charged residue of the Qλ arm tip that is predicted to be 5 Å from the negatively charged DNA-phosphate backbone by a positively charged residue, replacing an unfavorable electrostatic protein-DNA interaction by a favorable electrostatic interaction (Fig. S1B).

**Fig. 3.**
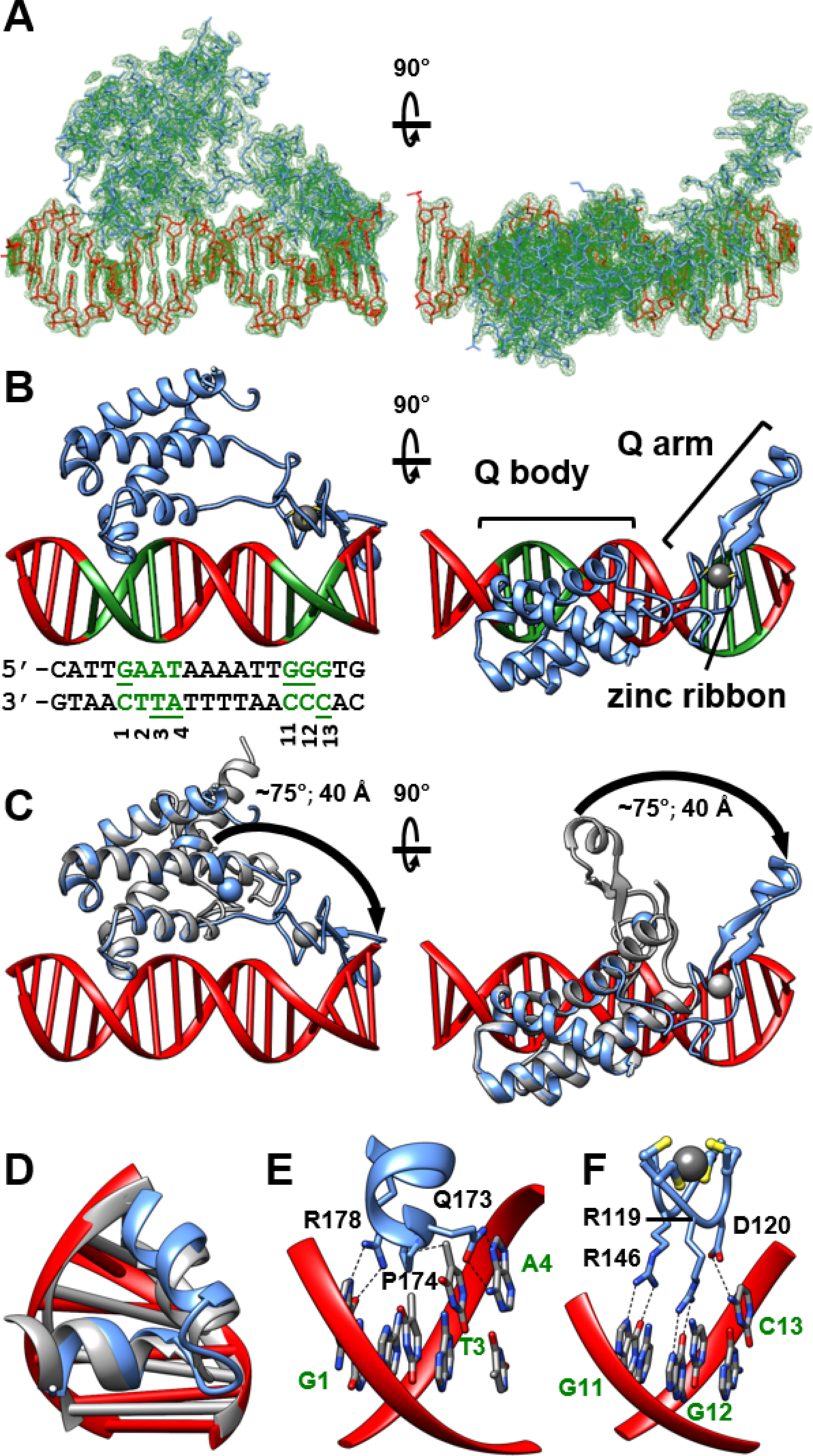
Structure of Q**λ**-QBE complex. **(A)** Structure of Qλ-QBE complex (two orthogonal views). Experimental electron density map (green mesh, 2mFo-DFc map contoured at 1.0σ) and fitted atomic model (red and blue). **(B)** Structure of Qλ-QBE complex (ribbon representation; two orthogonal views). Red and green, QBE DNA fragment (nucleotide pairs 1-4 and 11-13 in green; contacted bases underlined). Other colors as in Fig. 2B. **(C)** Superimposition of structure of Qλ (type-I crystal structure; gray ribbon; blue sphere for associated Zn^2+^) on structure of Qλ-QBE complex (blue and red ribbons for Qλ and QBE; gray sphere for associated Zn^2+^). **(D)** Qλ HTH motif (blue) interacting with DNA (red) superimposed on λ Cro HTH motif interacting with DNA (PDB 6CRO; gray). **(E)** Interactions of Qλ body with DNA major groove of nucleotide pairs 1-4. **(F)** Interactions of Qλ arm with DNA major groove of nucleotide pairs 11-13.

Comparison of the structures of Qλ-QBE and Qλ shows that, in Qλ-QBE, the Qλ arm is rotated, through a ∼75° swinging motion, resulting in a substantially more extended conformation (∼40 Å more extended for the Qλ arm tip) able to interact with a DNA site that spans more than one turn of DNA (Fig. 3C, cyan vs. gray ribbons). The structures suggest that formation of Qλ-QBE may involve two steps: a first step in which the Qλ body interacts with positions 1-4 of the QBE, and a second step in which the Qλ arm rotates, through a ∼75° swinging motion, to interact with positions 11-13 of the DNA site (Movie S1).

The structure of the Qλ-QBE complex, in which Qλ interacts as an extended monomer with an non-repeat, asymmetric DNA site (Fig. 3A-C), is radically different from the structure of the Q21-QBE complex, in which Q21 interacts as a dimer with a direct-repeat DNA site (12, 16). The results provide further evidence that Qλ and Q21 are not structural homologs.

### Structure of “pre-engaged” Qλ-loading complex

We have determined a single-particle-reconstruction cryo-EM structure of a Qλ-loading complex at 3.13 Å resolution (Figs. 4, S2-S3; Table S2). We prepared the Qλ-loading complex by *in vitro* reconstitution from ful-length Qλ, *Escherichia coli* RNAP core, an *E. coli* σ^70^ derivative having substitutions that increase efficiency of Qλ loading [R541C and L607P; (21–23)], and a nucleic-acid scaffold containing the λPR′ QBE, a consensus version of the λPR′ SDPE, and an 11 nt RNA (Fig. 4A). The DNA duplex contained a 16 bp non-complementary region overlapping the SDPE, corresponding to the unwound and scrunched transcription bubble in the Qλ-free, σ-containing paused TEC (pTEC) at λPR′ (24).

**Fig. 4.**
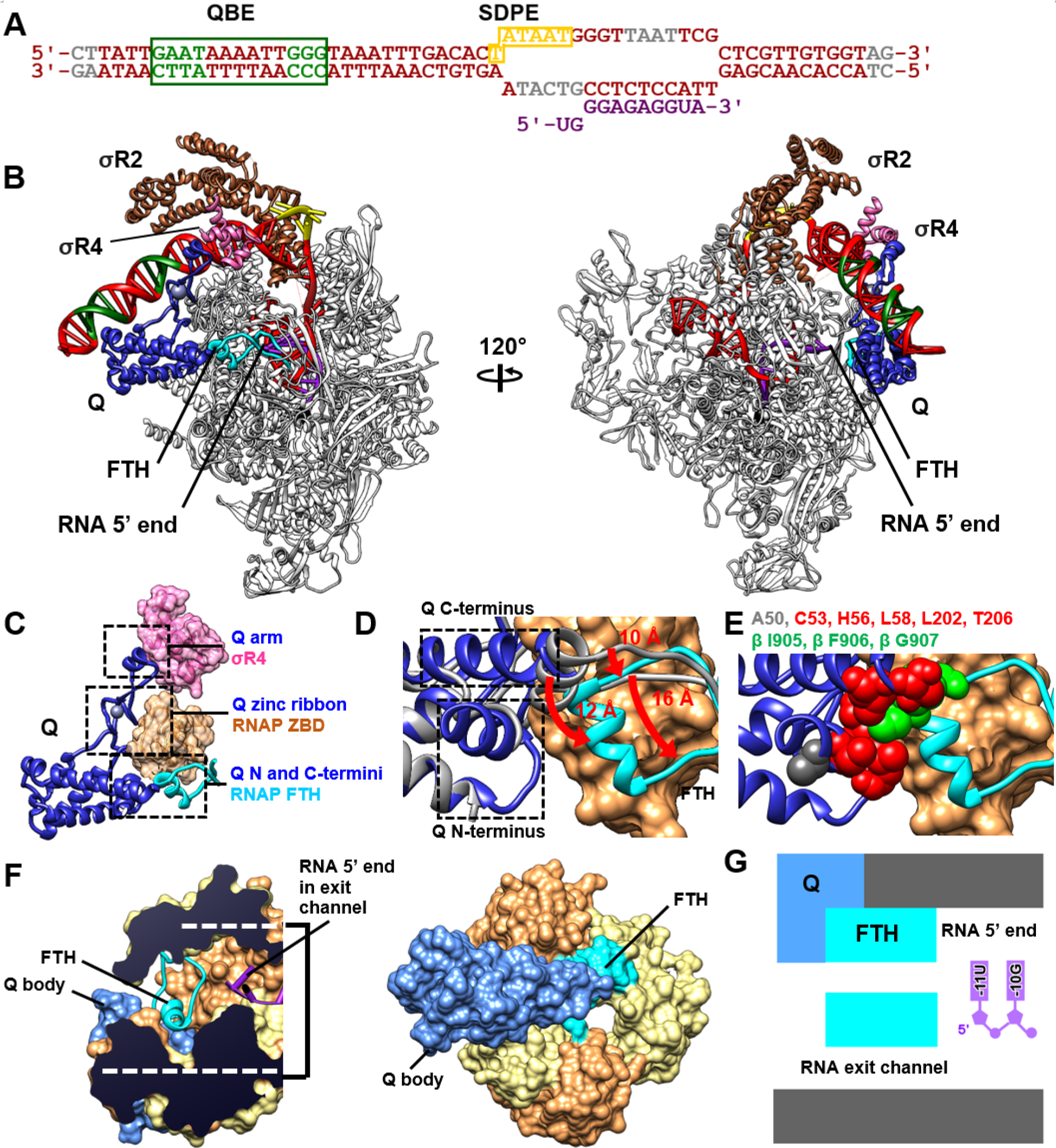
Structure of “pre-engaged” Qλ-loading complex. **(A)** Nucleic-acid scaffold. DNA, red (QBE nucleotide pairs 1-4 and 11-13, SDPE, and disordered nucleotides in green, yellow, and gray, respectively; non-complementary region corresponding to unwound transcription bubble indicated by raised and lowered letters); RNA, magenta. **(B)** Structure of pre-engaged Qλ-loading complex (two view orientations). Qλ, blue; RNAP, gray; RNAP FTH and connecting segments, cyan; σR2, brown; σR4, pink; DNA and RNA, colored as in A; RNAP active-center Mg^2+^, black sphere. **(C)** Qλ-RNAP interactions. Qλ, blue; RNAP ZBD, salmon. Other colors and view orientation as in left subpanel of B. **(D)** Conformational changes in Qλ upon formation of pre-engaged Qλ-loading complex. Qλ in Qλ-QBE complex (Fig. 3B, left; gray) superimposed on Qλ in pre-engaged Qλ-loading complex (colored as in B-C). Dashed rectangles, Qλ N-terminal residues that fold and C-terminal residues that refold upon formation of pre-engaged Qλ-loading complex (residues 46-60 and 195-207). **(E)** Interactions of Qλ N- and C-terminal segments with RNAP FTH in pre-engaged Qλ-loading complex. Sites of substitutions of Qλ that result in defects in Qλ-dependent antitermination but not Qλ-QBE interaction (35) are shown in red (residues that interact with FTH; C53, H56, L58, L202, and T206) and gray (residue that does not interact with FTH; A50). Sites of substitutions of FTH that result in defects in Qλ-dependent antitermination and Qλ-FTH interaction (23) are shown in green (I905, F906, and G907). View orientation as in D. **(F)** Qλ (blue) outside RNAP RNA-exit channel, RNAP FTH (cyan) partly in RNAP RNA-exit channel, and RNA (magenta; numbered assigning RNA 3’ nucleotide as -1) in RNAP RNA-exit channel. RNAP β and β’ are in salmon and light yellow, respectively. View orientation in left subpanel is orthogonal to RNA-exit channel; view orientation in right subpanel is parallel to RNA-exit channel. **(G)** Summary of organization of Qλ (blue), RNAP FTH (cyan), RNAP RNA-exit channel (gray), and RNA (magenta). View orientation as in left subpanel of F.

The structure shows Qλ interacting with the QBE in a manner matching that in the crystal structure of Qλ-QBE (Fig. 4B vs. Fig. 3A) and simultaneously interacting with a σ-containing paused TEC (Fig. 4B). The structural module of σ that recognizes the promoter -10 element in a transcription initiation complex, σ region 2 (σR2), makes interactions with the SDPE -10-element-like sequence and RNAP equivalent to those it makes in a transcription initiation complex (Fig. 4B). In contrast, the structural module of σ that recognizes a promoter -35 element in a transcription initiation complex, σ region 4 (σR4), makes interactions radically different from those it makes in a transcription initiation complex. In a transcription initiation complex, σR4 interacts with the RNAP flap-tip helix [FTH; β residues 897-907; (25–28)] and interacts with a -35 element ∼17 bp upstream of σR2 bound to a -10 element (27, 29–31). In the Qλ-containing pTEC, σR4 is disengaged from the RNAP FTH and is repositioned to a -35-element-like DNA site immediately upstream of σR2 bound to the SDPE -10-element-like sequence, where it makes protein-DNA interactions essentially identical to those it makes with a promoter -35 element in a transcription initiation complex (Figs. 4B-C, S4), consistent with published genetic and biochemical data (22, 32–34). The repositioned σR4 makes protein-protein interactions with Qλ (with the N-terminal part of σR4, residues 552-554, interacting with the Qλ arm tip, residues 133-137; Figs. 4B-C, S4), consistent with genetic and biochemical data indicating the importance of Qλ residue 134 and σ residue 553 for Qλ-σR4 interaction [Figs. 4B-C, S4; (17, 33)], and also makes unanticipated protein-protein interactions with σR2 (with the C-terminal part of σR4, residues 612-613, interacting with the σR2 non-conserved region, residues 154-155; Figs. 4B-C, S4). The structural modules of σ between σR2 and σR4, σ region 3 (σR3) and the σR3-σR4 linker, are disengaged from their positions in a transcription initiation complex--where σR3 interacts with DNA immediately upstream of σR2 bound to a -10 element, and where the σR3-σR4 linker occupies the RNAP RNA-exit channel (27, 30, 31)--and are disordered (Fig. 4B). Clear, traceable density is present for the entire 11 nt RNA. The RNA 5′-end nucleotide and one additional RNA nucleotide (positions -11 and -10) are in the RNAP RNA-exit channel, and the other nine RNA nucleotides (position -9 through -1) are base-paired to the transcription-bubble template DNA strand as an RNA-DNA hybrid (Fig. 4B).

Comparison of the structure of the Qλ-containing pTEC at λPR’ (Fig. 4) to the structure of the Qλ-free pTEC at λPR’ (24), reveals four structural changes induced by Qλ: (i) repositioning of σR4 to the DNA segment immediately upstream of σR2, (ii) displacement of σR3 from the DNA segment immediately upstream of σR2, (iii) bending of upstream DNA by ∼40° toward the RNAP ZBD, and (iv) repositioning of the RNAP FTH and the protein segments that precede and follow it, by 10-16 Å into the RNAP RNA-exit channel. The first of these structural changes is induced directly by Qλ, through direct Qλ-σR4 interaction (Figs. 4B-C, S4). The second is induced indirectly, by the repositioning of σR4 to the DNA segment immediately upstream of σR4 (Fig. 4B). The third and fourth are induced directly, through direct Qλ-ZBD and Qλ-FTH interactions (Fig. 4B-E).

Comparison of the structure of the Qλ-containing pTEC (Fig. 4) to the structure of the Qλ-QBE (Fig. 3), reveals that, upon interaction with the pTEC: (i) 16 residues of the 60-residue intrinsically disordered N-terminal segment of Qλ undergo a disorder-to-order transition, folding as a turn followed by an α-helix (residues 45-60; Fig. 4D; Movie S2), and (ii) 12 residues of the C-terminal segment of Qλ refold, extending the C-terminal α-helix of Qλ by three turns (residues 195-207; Fig. 4D; Movie S2).

In addition to interacting with, and re-positioning, σR4 (Figs. 4B-C, S4), Qλ interacts with RNAP in the Qλ-containing pTEC (Fig. 4B-E). The Qλ zinc ribbon (residues 110-112, 117, and 122-126) interacts with the RNAP zinc-binding domain (ZBD; β’ residues 75, 82-86, and 91; Fig. 4C), and the Qλ N-terminal and C-terminal regions (residues 52, 56, 58, 202, and 206) interact with the C-terminal half of the RNAP FTH (β residues 905-906; Fig. 4E). There is a nearly one-for-one correspondence between Qλ residues observed to make Qλ-FTH interactions in the structure and Qλ residues shown experimentally to be important for Qλ-dependent antitermination but not for Qλ-QBE interaction [Fig. 4E, red; (35)], suggesting that the observed interactions are functionally relevant. Furthermore, there is a one-for-one correspondence between RNAP FTH residues observed to make Qλ-FTH interactions in the structure and FTH residues previously shown experimentally to be important for Qλ-dependent antitermination and Qλ-FTH interaction [Fig. 4E, green; (23)], further suggesting that the observed interactions are functionally relevant.

In the Qλ-containing pTEC, Qλ-FTH interactions re-position part of the RNAP FTH into the RNAP RNA-exit channel (Fig. 4F-G). However, in contrast to the structure of the Q21 loading complex, in which Q21 forms a torus at the mouth of, and inside of, the RNAP RNA-exit channel that narrows and extends the RNAP RNA-exit channel (12, 16) in the structure of this Qλ-containing pTEC, Qλ does not interact with, or enter, the RNAP RNA-exit channel (Fig. 4F-G).

We term the structural state in Fig. 4, the “Qλ ‘pre-engaged’ loading complex,” to reflect the fact that this structural state has Qλ recruited to the pTEC, but does not have Qλ interacting with, or entering, the RNAP RNA-exit channel.

### Structure of NusA-containing “engaged” Qλ-loading complex

Qλ requires the transcription elongation factor NusA (10, 36, 37) for efficient antitermination (2, 4). NusA stabilizes the Qλ-loading complex (4) and increases the antitermination activity of the Qλ-loaded complex (2, 4).

We have determined a single-particle-reconstruction cryo-EM structure of a NusA-containing Qλ-loading complex, prepared as in the preceding section, but in the presence of NusA, at 3.36 Å resolution (Figs. 5, S5-S6; Table S2).

**Fig. 5.**
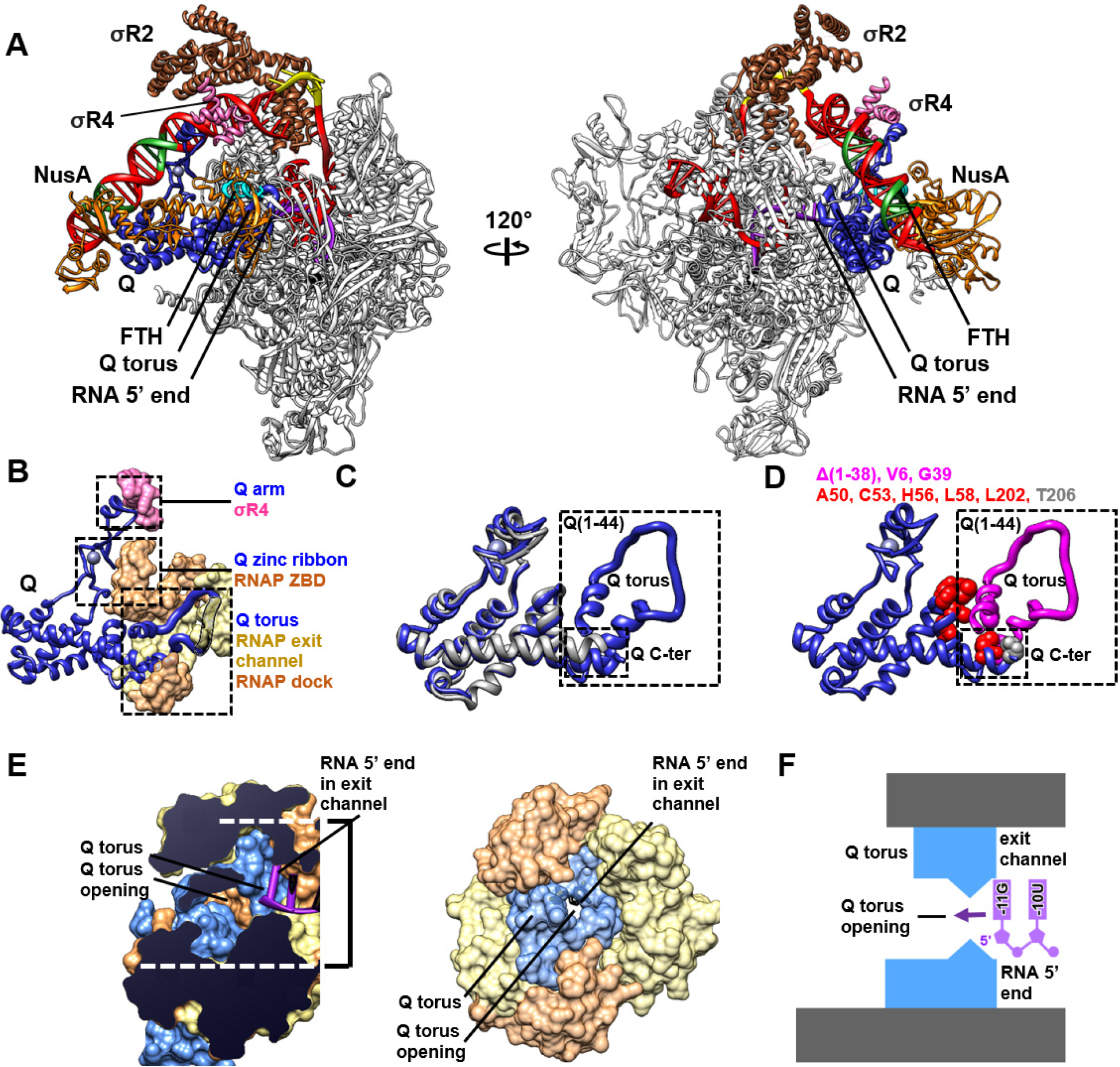
Structure of NusA-containing, “engaged” Qλ-loading complex. **(A)** Structure of NusA-containing, engaged Qλ-loading complex. NusA, orange. View orientations and other colors as in Fig. 4B. **(B)** Qλ-RNAP interactions. Qλ, blue, with segments located inside RNAP RNA-exit channel, behind RNAP surfaces, indicated as gray ribbons with black outlines. RNAP β and β’, salmon and light yellow, respectively. NusA omitted for clarity. Other colors and view orientation as in Fig. 4C. **(C)** Conformational changes in Qλ upon formation of NusA-containing, engaged Qλ-loading complex. Qλ in pre-engaged Qλ-loading complex (Fig. 4B, left; gray) superimposed on Qλ in NusA-containing, engaged Qλ-loading complex (colored as in A-B). Large dashed rectangle, Qλ N-terminal residues that fold to form Qλ torus (residues 1-44). Small dashed rectangle, Qλ C-terminal residues that interact with N-terminal region of Qλ torus. **(D)** Formation of Qλ torus at and inside RNAP RNA-exit channel in NusA-containing, engaged Qλ-loading complex. Qλ residues deleted in mutant defective in Qλ-dependent antitermination, but not defective in Qλ-QBE interaction [residues 1-38; (17)], and Qλ residues altered in single-substitution mutants defective in Qλ-dependent antitermination, but not defective in Qλ-QBE interaction [V6A and G39A; (35)], are shown as pink ribbons and pink surfaces, respectively. Sites of substitutions of Qλ that result in defects both in Qλ-dependent antitermination but not Qλ-QBE interaction (35) are shown in red (residues that interact with Qλ torus; C53, H56, L58, L202, and T206) and gray (residue that does not interact with Qλ torus; T206). View orientation and dashed rectangles as in C. **(E)** Restriction of RNAP RNA-exit channel by Qλ torus (blue) and proximity of RNA 5’ end (magenta; RNA nucleotides numbered assigning RNA 3’ nucleotide as -1) to Qλ torus. RNAP β and β’ are in salmon and light yellow, respectively. View orientation in left subpanel is orthogonal to RNA-exit channel; view orientation in right subpanel is parallel to RNA-exit channel. **(F)** Summary of organization of Qλ torus (blue), RNAP RNA-exit channel (gray), and RNA (magenta). View orientation as in left subpanel of E.

The NusA-containing Qλ-loading complex (Fig. 5) has the same overall structural organization as the NusA-free, pre-engaged Qλ-loading complex (Fig. 4). Qλ interacts with the QBE through the Qλ body and Qλ arm and simultaneously interacts with a σ-containing pTEC, with the Qλ body interacting with the RNAP ZBD and the Qλ arm tip interacting with σR4 re-positioned to the DNA segment immediately upstream of σR2 bound to the SDPE -10-like element (Fig. 5A-B).

In the NusA-containing Qλ-loading complex, NusA interacts with both RNAP and Qλ (Figs. 5A, S7). The NusA N-terminal domain interacts with the RNAP FTH and with one RNAP α-subunit C-terminal domain, making interactions equivalent to those it makes in other structures of NusA-containing TECs [Figs. 5A, S7; (38–44)]. The NusA S1 domain (residues 144-147 and 170-175) interacts with the Qλ body (residues 80-81, 106-107, and 182-197) (Figs. 5A, S7).

Comparison of the structure of the NusA-containing Qλ-loading complex (Fig. 5) to the structure of the NusA-free, pre-engaged Qλ-loading complex (Fig. 4) shows that, upon binding of NusA, remarkable large-scale conformational changes occur in the RNAP FTH and in Qλ. The RNAP FTH, which interacts with Qλ in NusA-free, pre-engaged Qλ-loading complex (Fig. 4), moves by 26 Å, breaking its interactions with Qλ, and, instead, making interactions with the NusA N-terminal domain (Figs. 5A, S8). Residues 1-44 of Qλ, which are disordered in Qλ, in Qλ-QBE, and in the NusA-free, pre-engaged Qλ-loading complex (Figs. 3-4), undergo a disorder-to-order transition to form a Q torus and to position the Q torus at the mouth of, and inside, the RNAP RNA-exit channel, in space vacated by the re-positioned RNAP FTH (Fig. 5; Movie S3). Residues 1-44 of Qλ fold to form a first α-helix, followed by a loop, followed by a second α-helix (residues 1-11, 12-31, and 32-42, respectively). The first α-helix interacts with residues of the RNAP ZBD at the mouth of the RNA-exit channel (β’ residues 67, 78-79, 94, and 96), the loop interacts with residues of the RNAP ZBD (β’ residue 49) and RNAP flap (β residues 843-844, 848, 888, 914-917, and 919) inside the RNA-exit channel, and the second α-helix interacts with residues of the RNAP dock (β’ residues 386, 394, 397-398) and RNAP β subunit (β residues 1302 and 1305-1306) at the mouth of the RNA-exit channel (Fig. 5A-D).

There is an almost one-for-one correlation between the Qλ residues that undergo the disorder-to-order transition to form the Qλ torus upon NusA binding (residues 1-44) and the Qλ residues deleted in a deletion mutant defective in Qλ-dependent antitermination but not defective in Qλ-QBE and Qλ-RNAP interactions [residues 1-38; (17); Fig. 5D, pink ribbon]. There also is a correlation between Qλ residues that form the Qλ torus and Qλ residues altered in single-substitution mutants defective in Qλ-dependent antitermination but not defective in Qλ-QBE interaction [V6A and G39A; (35); Fig. 5D, pink surfaces]. We conclude that the Qλ torus is functionally significant for Qλ-dependent antitermination.

The binding of the Q torus at the mouth of, and inside, the RNAP RNA-exit channel markedly restricts the RNAP RNA-exit channel, narrowing and extending the channel (Fig. 5E-F). The Q-torus opening has a solvent-excluded diameter of just 5-7 Å (Fig. 5E). The presence of the Q torus at, and inside, the mouth of the RNAP RNA-exit channel does not affect accommodation of an 11-nt RNA product (Fig. 5A,F), but would necessitate longer RNA products to thread through the Q-torus opening.

The Qλ torus in the structural state in Fig. 5 has no sequence, secondary-structure, or tertiary-structure similarity to the Q21 torus in structures of the Q21-loading complex and Q21-loaded complex (12, 16). However, the Qλ torus opening has the same diameter as the Q21 torus opening [solvent-excluded diameter of 5-7 Å; Fig. 5E; (12, 16)], the Qλ torus extends the same distance into the RNAP RNA-exit channel as the Q21 torus [accommodating a maximum of 11-13 nt of RNA before threading of RNA through the torus; Fig. 5E-F; (12)], and the Qλ torus has the same high net positive charge as the Q21 torus, enabling threading of RNA through interactions solely with the negatively charged RNA phosphate backbone [net positive charge of +4, excluding side chains that interact with RNAP; (12)]. Thus, in the structural state in Fig. 5, Qλ forms a nozzle at, and inside, the RNAP RNA-exit channel that is not homologous to, but that is analogous in all functionally important respects to, the nozzle previously observed in the Q21-loading complex and Q21-loaded complex.

We term the structural state in Fig. 5, the “Qλ ‘engaged’ loading complex,” to reflect the fact that this structural state not only has Qλ recruited to the pTEC, but also has Qλ reorganized to form a molecular nozzle interacting with, and entering, the RNAP RNA-exit channel.

## DISCUSSION

### Qλ: mechanism of loading onto RNAP

The functional importance of the interactions between Qλ and the RNAP FTH that are observed in our structure of the pre-engaged Qλ-loading complex, but that are not observed in our structure of the NusA-containing, engaged Qλ-loading complex, is validated by the finding that substitution of Qλ and FTH residues that make those interactions results in a specific defect in Qλ-dependent antitermination [Fig. 4E, red and green; (23, 35)] and by the further finding that substitution of FTH residues that make those interactions results in a specific defect in Qλ-FTH interaction [Fig. 4E, green; (23)]. The functional importance of the interactions between the Qλ torus and the RNAP RNA-exit channel that are observed in our structure of the NusA-containing, engaged Qλ-loading complex, but that are not observed in our structure of the pre-engaged Qλ-loading complex, is validated by the finding that deletion or substitution of residues that make those interactions results in a specific defect in Qλ-dependent antitermination [Fig. 5D, pink ribbon and pink surfaces; (17, 35)]. We conclude that *both* the pre-engaged Qλ-loading complex and the NusA-containing, engaged Qλ-loading complex are *bona fide*, on-pathway intermediates in Qλ loading. We propose that Qλ loading involves two stages: (i) recruitment of Qλ, yielding a pre-engaged Qλ-loading complex, and (ii) reorganization of Qλ to form a Qλ torus and engage the RNAP RNA-exit channel, yielding an engaged Qλ-loading complex. We propose further that NusA facilitates the transition from the pre-engaged Qλ-loading complex to the engaged Qλ-loading complex.

Comparison of the structure of the pre-engaged Qλ-loading complex at λPR’ (Fig. 4) to the structure of a Qλ-free pTEC at λPR’ (24), together with consideration of functional data for Qλ-RNAP interaction (21, 23, 35), indicates that formation of the pre-engaged Qλ-loading complex from the Qλ-free pTEC involves the following events: (i) binding of Qλ to the QBE; (ii) repositioning of σR4 to the DNA segment upstream of σR2 bound to the SDPE -10-like element, displacing σR3 from the DNA segment, (iii) bending of the DNA segment between the QBE and the pTEC by ∼40°, enabling interaction of the Qλ body with the RNAP ZBD; (iii) folding of 16 residues at the Qλ N-terminus and refolding of 12 residues at the Qλ C-terminus; and (iv) interaction of the Qλ N-and C-terminal segments with the RNAP FTH (Fig. 6A, left).

**Fig. 6.**
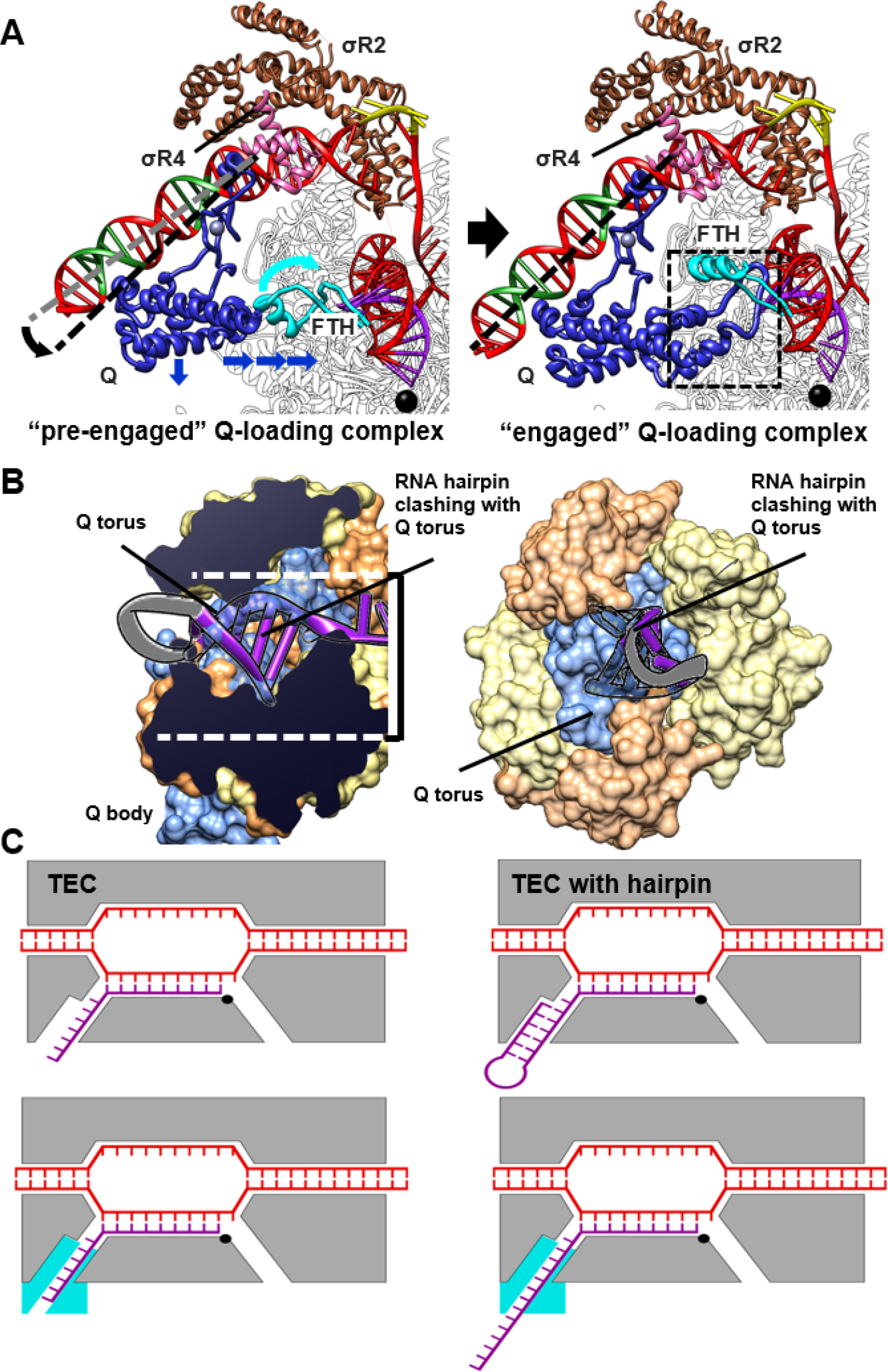
Mechanistic conclusions. **(A)** Transformation of pre-engaged Qλ-loading complex to NusA-containing, engaged Qλ-loading complex (NusA not shown for clarity). Black and gray dashed lines, DNA-helix axes of upstream dsDNA segments in pre-engaged and engaged complexes, respectively; blue arrows, cyan arrow, and dashed rectangle, structural changes upon transformation of pre-engaged to engaged complex. Other colors as in Figs. 4B and 5A. **(B)** Antitermination and antipausing by Qλ. Top, steric incompatibility of Qλ torus with pause and terminator RNA hairpins (hairpin stem from PDB 6ASX in purple, with segments positioned to interpenetrate Qλ torus indicated as transparent ribbons with black outlines; hairpin loop from PDB 1MT4 as gray ribbon with black outlines). View orientations and colors as in Figs. 4F and 5E. **(C)** Schematic comparison of TECs in absence of Qλ (upper row) and TECs in presence of Qλ (lower row). Colors as in Fig. 4B and 5A.

Comparison of the structure of the NusA-containing, engaged Qλ-loading complex at λPR’ (Fig. 5) to the structure the pre-engaged Qλ-loading complex at λPR’ (Fig. 4), together with consideration of functional data for Qλ-dependent antitermination (17, 35), indicates that formation of the NusA-containing, engaged Qλ-loading complex from the pre-engaged Qλ-loading complex involves the following events: (i) binding of NusA to Qλ and RNAP, making interactions with the Qλ body and RNAP α C-terminal domain, stabilizing the association between Qλ and RNAP; (ii) interaction of the NusA N-terminal domain with the RNAP FTH, disrupting interactions between Qλ and the FTH, and repositioning the FTH outside of and away from the RNAP RNA-exit channel; and (iii) folding of 44 additional residues at the Qλ N-terminus to form a Qλ torus that enters the RNAP RNA-exit channel (Fig. 6A, right).

The inferred events in Qλ loading include a remarkable “two-handoff mechanism.” In formation of the pre-engaged Qλ-loading complex, the first handoff occurs: i.e., the RNAP FTH is “handed off” from σR4--which interacts with the RNAP FTH in the absence of Qλ (25-28, 30, 31, 45)--to Qλ. In formation of the NusA-containing, engaged Qλ-loading complex, the second handoff occurs: i.e., the FTH is “handed off” from Qλ to NusA.

Whereas Qλ employs a NusA-dependent, two-stage process for Q loading--involving recruitment of Q to form a pre-engaged loading complex, followed by NusA-facilitated refolding of Q to form an engaged loading complex with a Q torus in the RNAP RNA exit channel--Q21 loading occurs in a NusA-independent, single-stage process (12, 16). Nevertheless, Q21 loading shows a fundamental underlying analogy to Qλ loading. The Q21-QBE complex and Q21 loading complex each contains two Q21 protomers, Q21u and Q21d, where “u” denotes “upstream” and “d” denotes “downstream” (12, 16). Q21u makes interactions analogous to those made by Qλ in the NusA-containing, engaged Qλ-loading complex, interacting with the RNAP ZBD and forming a Q torus that enters the RNAP RNA-exit channel. Q21d makes interactions analogous to those made by NusA in the NusA-containing, engaged Qλ-loading complex, interacting with Q21u and interacting with, and repositioning, the RNAP FTH. Thus, together, Q21u and Q21d make interactions analogous to the key Qλ-RNAP, Qλ-NusA, and NusA-RNAP interactions in the NusA-containing, Qλ-loading complex.

### Qλ: mechanisms of antitermination and antipausing

The structure of the NusA-containing, engaged Qλ-loading complex immediately suggests the mechanisms of antipausing and antitermination by Qλ (Figs. 5E-F, 6B-C). RNA-hairpin-dependent transcription pausing and transcription termination involve nucleation of an RNA hairpin at the mouth of the RNAP RNA-exit channel, followed by propagation of the RNA hairpin stem and penetration of the RNA hairpin stem into the RNAP RNA-exit channel (46, 47). The structure of the NusA-containing, engaged Qλ-loading complex shows that Qλ is positioned to pose a steric barrier to nucleation, propagation, and penetration of the RNA-exit channel by a pause or terminator hairpin (Fig. 6B-C). The Qλ torus is positioned to overlap, *in toto*, 4-5 bp of the stem of a pause or terminator hairpin, and thus is positioned to block propagation and penetration of the RNA-exit channel by a hairpin (Fig. 6B-C).

Because the Qλ torus has dimensions that accommodate only single-stranded RNA, the Qλ torus constitutes an effectively absolute steric barrier to the formation of double-stranded RNA secondary structure at the mouth of, or inside, the RNAP RNA-exit channel (Fig. 6B-C).

The structure of the NusA-containing, engaged Qλ-loading complex suggests that Qλ also may exert antipausing activity by inhibiting RNAP swiveling--a rotation, by ∼3°, of the RNAP “swivel module,” comprising the RNAP ZBD and associated RNAP domains, that has been proposed to be associated with pausing [Fig. S8; (38, 39, 48–50)]. In the structure of the NusA-containing, engaged Qλ-loading complex, the RNAP swivel module is in the unswivelled state (Fig. S8A), despite the presence of NusA, which favors the swivelled state (44). Model building indicates that interaction of the Qλ body with one face of the RNAP ZBD and the interaction of an α-helix of the Qλ torus with the other face of the RNAP ZBD would sterically preclude both the ∼3° swiveling associated with hairpin-dependent pausing (38, 39) and NusA binding (44) and the smaller, ∼1.5°, swiveling associated with elemental pausing [(38); Fig. S8].

The structure of the NusA-containing, engaged Qλ-loading complex implies that, upon further RNA extension, RNA would thread into, and through, the Qλ torus (Figs. 5E-F, 6B-C). Because re-threading of the extruded RNA would be difficult or impossible, especially after RNA folding, the threading of RNA through the Qλ torus would result in a topological linkage between RNA and Qλ, creating an essentially unbreakable linkage between the TEC and Qλ, enabling processive antitermination and antipausing over tens of thousands of nucleotide-addition steps (11).

The structures of this work indicate that the Qλ torus, like the Q21 torus (12, 16), forms a molecular nozzle, that narrows and extends the RNAP RNA-exit channel, preventing the formation of pause and terminator hairpins, and that blocks formation of the RNAP swiveled state associated with pausing (Figs. 6C, S8B).

### Qλ and Q21: functional analogy without structural homology

Our results show that Qλ exhibits no three-dimensional structural similarity to Q21 [Figs. 2-5; (12, 16)], show that Qλ employs a different mechanism for DNA binding than Q21 [Fig. 3; (12, 16)], and show that Qλ employs a different, more complex, process of loading onto TECs than Q21 [Figs. 4-6; (12, 16)]. Our results indicate, nevertheless, that Qλ employs the same molecular-nozzle mechanism for antipausing and antitermination as Q21 [Figs. 5-6; (12, 16)]. We conclude that Qλ and Q21 are not structural homologs and are solely functional analogs.

### Prospect

One priority for further research is to determine whether the third protein family of lambdoid bacteriophage Q proteins, the Q82 protein family, also functions through a nozzle mechanism. The observation that Q82 can load onto a TEC that contain a long RNA product (51)--which should be topologically difficult or impossible for a factor functioning as a closed molecular nozzle--raises the possibility that Q82 might not function as a closed molecular nozzle.

Another priority for further research emerges from the hypothesis that nozzle formation by Q inhibits termination by extending the RNAP RNA-exit channel. The hypothesis implies that the same nozzle formation that prevents termination at standard terminators that contain a hairpin immediately followed by a 7-9 nt U-tract (46, 47), potentially could induce termination at non-standard, “Q-dependent” terminators that contain a hairpin, followed by a spacer--an extension matching the extension of the RNAP RNA-exit channel--followed by a 7-9 nt U-tract [see discussion in (7)]. In principle, such non-standard, Q-dependent terminators could have functional roles in ending processive Q-dependent antitermination. Construction and analysis of libraries of terminator derivatives containing spacers between the hairpin and the U-tract, combined with analysis of corresponding sequences in bacteriophage or bacterial genomes, could address whether such non-standard, Q-dependent terminators exist, and, if so, whether they play functional roles.

## ACKNOWLEDGEMENTS

This work was supported by National Institutes of Health grant GM118059 (to B.E.N.) and GM041376 (to R.H.E.). We thank the Argonne National Laboratory for beamline access, the Rutgers Cryo-EM and Nanoimaging Facility and the National Center for CryoEM Access and Training (supported by NIH grant GM129539, Simons Foundation Grant SF349247, and New York state grants) for microscope access, and J. Roberts for plasmids.

## SUPPORTING INFORMATION APPENDIX

### SUPPLEMENTARY MATERIALS AND METHODS

#### Bacteriophage λ Q derivatives

Qλ^M1S^ and Qλ^61-207;N61S;E134K^ were prepared from cultures of *E. coli* strain BL21Star(DE3) (Invitrogen, Inc.) transformed with plasmid pET21-H6-TEV-Qλ^M1S^ or pET21-H6-TEV-Qλ^61-207;N61S;E134K^, encoding N-terminally hexahistidine-tagged, tobacco-etch-virus-protease-site-tagged Qλ^M1S^ or Qλ Qλ^61-207;N61S;E134K^ under the control of the bacteriophage T7 gene 10 promoter [constructed by replacing the NdeI-SalI segment of plasmid pET21a (EMD Millipore, Inc.) with an NdeI-SalI segment of a synthetic DNA fragment (Genscript, Inc.) carrying 5’-CATATGGGACATCACCATCACCATC ACGAGAACCTGTACTTCCAATCC-3’, followed by codons 2-207 or codons 62-207 with E134K mutation (1) of gene Q of lambdoid bacteriophage λ (NCBI NP_040642.1), followed by 5’-TGAGTCGAC-3’]. Protein production, protein purification, and protein tag removal were performed as for 21Q derivatives in (2), and proteins were concentrated to 20 mg/ml in buffer A (10 mM Tris-HCl, pH 7.4, 100 mM NaCl, and 1 mM dithiothreitol) using 10 kDa MWCO Amicon Ultra-15 centrifugal ultrafilters (Millipore, Inc.) and stored at -80°C. Yields were ∼5 mg/L, and purities were >95%.

Wild-type Qλ was prepared from cultures of *E. coli* strain BL21(DE3) (Invitrogen, Inc.) transformed with plasmid pQE30-λQ (3), as in (4).

*E. coli* RNAP core enzyme *E. coli* RNAP core enzyme containing C-terminally decahistidine-tagged β’ subunit was prepared from cultures of *E. coli* strain BL21(DE3) (Invitrogen, Inc.) transformed with plasmid pIA900 [encodes *E. coli* RNAP β’, β, α, and ω subunits under control of the bacteriophage T7 gene 10 promoter; (5)], as in (5).

*E. coli* [Cys541;Pro607]-σ^70^ N-terminally hexahistidine-tagged [Cys541;Pro607]-σ^70^ [*E. coli* σ^70^ derivative having reduced affinity for the RNAP FTH, resulting in increased proficiency in Q loading; (6–8)] was prepared from cultures of *E. coli* strain BL21Star(DE3) (Invitrogen, Inc.) transformed with plasmid pLHN12-His-R541C;L607P (6), as in (9).

E. coli NusA N-terminally hexahistidine-tagged NusA was prepared from cultures of *E. coli* strain BL21Star(DE3) (Invitrogen, Inc.) transformed with pET15b-NusA (10), as in (10).

#### Oligonucleotides

Oligodeoxyribonucleotides (IDT), and oligoribonucleotides (IDT) were dissolved in nuclease-free water (Ambion, Inc.) to 1 mM and stored at -80°C.

#### Nucleic-acid scaffolds

The nucleic-acid scaffold for preparation and analysis of the Qλ-QBE complex (sequence in Figs. 3B and S1) was prepared as follows: Nontemplate-strand oligodeoxyribonucleotide (0.5 mM), and template-strand oligodeoxyribonucleotide (0.5 mM) in 500 µl 10 mM Tris-HCl, pH 8.0, and 50 mM NaCl were heated 10 min at 95°C, slow-cooled to 25°C (∼2 h), and stored at -80°C.

The nucleic-acid scaffold for preparation and analysis of the pre-engaged Qλ-loading complex and the NusA-containing, engaged Qλ-loading complex (sequence in Fig. 4A) was prepared as follows: Nontemplate-strand oligodeoxyribonucleotide (0.3 mM), template-strand oligodeoxyribonucleotide (0.3 mM), and oligoribonucleotide (0.33 mM) in 500 µl 10 mM Tris-HCl, pH 8.0, and 50 mM NaCl were heated to 95°C for 2.5 min, slow-cooled to 25°C (∼3 h) and then to 4°C (∼0.5 h), and stored at -80°C.

#### Crystal-structure determination: sample preparation

The Qλ^61-207;N61S;E134K^-QBE complex was prepared by incubating 500 μM Qλ^61-207;N61S;E134K^ and 500 μM QBE nucleic-acid scaffold in 2 ml buffer A for 10 min at 25°C, was purified by size-exclusion chromatography on a Superdex S75 16/60 column (GE Healthcare, Inc) equilibrated and eluted in 10 mM MES pH 6.5, 50 mM NaCl, and 1 mM dithiothreitol, was concentrated to 6 mg/ml using 10 kDa MWCO Amicon Ultra-15 centrifugal ultrafilters (Millipore, Inc.), was flash frozen in liquid nitrogen, and was stored at -80°C.

#### Crystal-structure determination: crystallization and data collection

Robotic crystallization trials for Qλ^61-207;N61S;E134K^ (Qλ here and below) and Qλ^61-207;N61S;E134K^-QBE (Qλ-QBE here and below) were performed using a Gryphon liquid handling system (Art Robbins Instruments, Inc.), commercial screening solutions (Emerald Biosystems, Inc, Hampton Research, Inc., and Qiagen, Inc.), and the sitting-drop vapor diffusion technique [drop: 0.5 µl μM Qλ or Qλ-QBE in buffer C plus 0.5 µl screening solution; reservoir: 60 µl screening solution; 6°C or 18°C]. Nine hundred conditions each were screened for Qλ and Qλ -QBE, and conditions yielding crystals subsequently were optimized using the hanging-drop vapor-diffusion technique.

Type-I crystals of Qλ for X-ray diffraction data collection were prepared using the sitting-drop vapor-diffusion method, by mixing 0.5 µl 800 µM Qλ and Qλ -QBE in buffer C with 0.5 μl of a reservoir solution consisting of 20% (v/v) glycerol, 16% (w/v) polyethylene glycol 8000 (Hampton Research, Inc.), and 400 mM potassium phosphate dibasic at 18°C.

Type-II crystals of Qλ for X-ray diffraction data collection were prepared using the sitting-drop vapor-diffusion method, by mixing 0.5 µl 800 μM Qλ in buffer C with 0.5 μl of a reservoir solution consisting of 10% (v/v) glycerol, 30% (v/v) polyethylene glycol 600 (Hampton Research, Inc.), 100 mM

Hepes-NaOH, pH 7.5, and 50 mM lithium sulfate, at 18°C.

Crystals of Qλ-QBE for X-ray diffraction data collection were prepared using the hanging-drop vapor-diffusion method, by mixing 1 µl 230 µM Qλ-QBE in 10 mM MES, pH 6.5, 50 mM NaCl, and 1 mM dithiothreitol with 1 µl reservoir solution consisting of 1 M sodium citrate and 100 mM Hepes-NaOH, pH 7.0.

Crystals of Qλ were cryo-protected in their own reservoir solution and were cooled by immersion in liquid nitrogen. Crystals of Qλ-QBE were cryo-protected by transfer to 5 μl reservoir solution containing 15% ethylene glycol and were cooled by immersion in liquid nitrogen.

X-ray diffraction data were collected on single crystals of Qλ and Qλ-QBE at 100°K at the Argonne National Laboratory Advanced Photon Source beamline 19-ID (wavelength 0.987 Å).

#### Crystal-structure determination: structure solution and refinement

Data were processed, and integrated intensities were merged and scaled, with automatic correction and a resolution cut-off of I/σ = 2.0, using HKL2000 (11).

The type-I crystal structure of Qλ was solved by molecular replacement in Phenix (12), using the crystal structure of Qλ^39-207^ [PDB 4MO1, (13)] as the search model.

The type-II crystal structure of Qλ was solved by molecular replacement in Phenix (12), using the type-I crystal structure of Qλ (Fig. 2), minus the Qλ arm (residues 113-157), as the search model.

The crystal structure of Qλ-QBE was solved by experimental phasing with single-wavelength anomalous dispersion (14) in Phenix (12), using zinc anomalous diffraction data. One heavy-atom site was identified, and an initial model was built in Phenix.

Iterative cycles of model building in Coot (15) and refinement in Phenix (12)then were performed, adding riding hydrogen atoms to all models, performing translation-libration-screw-rotation parameterization by TLSMD analysis (16), and performing validation with MolProbity (17).

Final models and structural factors of the Qλ type-I crystal structure (protomer I residues 62-206 and protomer II residues 62-205) refined to 1.46 Å, the Qλ type-II crystal structure (protomer I residues 62-206 and protomer II residues 62-207) refined to 1.97 Å, and the Qλ-QBE crystal structure (Qλ residues 62-201, nontemplate-strand nucleotides 1-19, and template-strand nucleotides 1-19) refined to 2.18 Å were deposited in the Protein Data Bank (PDB) with accession codes 6VEU, 6VEV and 6VEW, respectively (Table S1).

#### Cryo-EM structure determination: sample preparation

The pre-engaged Qλ-loading complex was prepared by incubating 6 μM *E. coli* RNAP core enzyme with 10 μM nucleic-acid scaffold (sequence in Fig. 4A) in 1 ml buffer B (10 mM Tris-HCl, pH 7.8, 100 mM NaCl, 1 mM MgCl_2_, and 1 mM dithiothreitol) for 5 min at 25°C, followed by adding 300 μl 30 μM *E. coli* [C541;P607]-σ^70^ in buffer B and 60 μl 360 μM Qλ^M1S^ in buffer A, and incubating for 5 min at 25°C. The sample was purified by size-exclusion chromatography on a Superdex S200 26/60 column (GE Healthcare, Inc.) equilibrated and eluted in buffer B, was concentrated to 9 mg/ml in buffer B using 30 kDa MWCO Amicon Ultra-15 centrifugal ultrafilters (Millipore, Inc.), and was stored at -80°C.

The NusA-containing, engaged Qλ-loading complex was prepared by incubating 2.2 μM *E. coli* RNAP core enzyme with 3.3 μM nucleic-acid scaffold (sequence in Fig. 4A) in 1 ml buffer B for 5 min at 25°C, followed by adding 20 μl 170 μM *E. coli* [C541;P607]-σ^70^ in buffer B, 40 μl 130 μM wild-type Qλ and 30 μl 300 μM NusA in buffer B, and incubating for 5 min at 25°C. The sample was purified by size-exclusion chromatography on a Superdex S200 26/60 column (GE Healthcare, Inc.) equilibrated and eluted in buffer B, was concentrated to 9 mg/ml in buffer B using 30 kDa MWCO Amicon Ultra-15 centrifugal ultrafilters (Millipore, Inc.), and was stored at -80°C.

EM grids were prepared using a Vitrobot Mark IV autoplunger (FEI, Inc.), with the environmental chamber at 22°C and 100% relative humidity. Samples comprising 45 μl 20 μM pre-engaged Qλ-loading complex or NusA-containing, engaged Qλ-loading complex in buffer B were mixed with 5 μl 80 mM CHAPSO (Hampton Research, Inc.) in water for 5 min on ice. Aliquots (2.5 μl) were applied to glow-discharged (5 min) UltrAuFoil (1.2/1.3) 300-mesh grids (Quantifoil, Inc.), grids were blotted with #595 filter paper (Ted Pella, Inc.) for 2 s at 22°C, grids were flash-frozen by plunging in a liquid ethane cooled with liquid nitrogen, and grids were stored in liquid nitrogen.

#### Cryo-EM structure determination: data collection and data reduction: pre-engaged Qλ-loading complex

Cryo-EM data were collected at the National Center for CryoEM Training and Access at hte Simons Electron Microscopy Center, using a 300 kV Titan Kiosk electron microscope (FEI/ThermoFisher, Inc.) equipped with a K2 Summit direct electron detector (Gatan, Inc.). Data were collected automatically in super-resolution counting mode using Leginon (18), a nominal magnification of 22,500x (actual magnification 47,608x), a calibrated pixel size of 0.533 Å per super-resolution pixel, and a dose rate of 1.98 electrons/pixel/s. Movies were recorded at 200 ms/frame for 8 s (40 frames total), resulting in a total radiation dose of 56.2 electrons/Å^2^. Defocus range was varied between -0.80 µm and -1.5 µm. A total of 3,027 micrographs were recorded from one grid over two days. Micrographs were saved as 4-bit LZW compressed Tiff images.

Data processing was performed on a Tensor TS4 Linux GPU workstation (Exxact, Inc.) containing four GTX 1080 Ti graphic cards (Nvidia, Inc.). Dose weighting, motion correction (5x5 tiles; b-factor = 150), Fourier-binning (2x2; pixel size = 1.066 Å), gain normalization, and defect correction were performed using MotionCor2 (19). Contrast-transfer-function (CTF) estimation was performed using Gctf (20).

Subsequent image processing was performed using Relion 3.0 (21).

Data were processed as summarized in Fig. S2A. Automatic particle picking with Laplacian-of-Gaussian filtering yielded an initial set of 1,337,270 particles. Particles were extracted into 256x256 pixel boxes, 4x down-scaled, and subjected to rounds of reference-free 2D classification and removal of poorly populated classes, yielding a selected set of 322,949 particles. The selected set was 3D-classified with C1 symmetry, using a 60 Å low-pass-filtered map calculated from a cryo-EM structure of an *E. coli* TEC [PDB 6ALF; (22)] as the 3D template, and the best-resolved classes, comprising 213,805 particles, were 3D auto-refined and then subjected to 3D sub-classification without alignment, using the auto-refined map as the 3D template. Following sub-classification, sub-classes exhibiting resolved Qλ-QBE density were combined, and the resulting 63,014 particles were re-centered, re-extracted without down-scaling, and subjected to 3D auto-refinement without a reference mask, using the previous auto-refined map as the 3D template. The resulting refined particles were CTF refined, Bayesian polished (23), and further CTF refined. The resulting CTF-refined particles were refined, and then subjected to a focused classification without alignment, using a custom mask generated by cropping the density volume of Qλ-QBE in Chimera (24) and converting it to a soft-edged mask in Relion. From the focused classification, a sub-class exhibiting resolved density for Qλ-QBE was selected and subjected to a focused classification using a custom mask generated by cropping density volume near the RNAP FTH in Chimera and converting it to a soft-edged mask in Relion. A sub-class exhibiting resolved density for the RNAP FTH was selected (11,913 particles) and refined, yielding a reconstruction at 3.13 Å overall resolution, as determined from gold-standard Fourier shell correlation [FSC; (25, 26)], using phase-randomization to account for convolution effects of a solvent mask on FSC between two independently refined half maps, “half-map 1” and “half-map 2” [(27); Figs. S2-S3; Table S2].

The initial atomic model for the pre-engaged Qλ-loading complex was built by manual docking of the crystal structure of Qλ-QBE (Fig. 3), DNA segments, RNA segments, RNAP β’, β, α^I,^, α^II^, ω and σ^70^ segments from a cryo-EM structure of Q21-loading complex [PDB 6P18; PMID: (2)], and an σR4 segment from a crystal structure of an *E. coli* transcription initiation complex [PDB 4YLN; (28)], using UCSF Chimera (24). For the Qλ N-terminus (residues 2-44), the RNAP β’ N and C-termini and SI3 (residues 1-14, 1376-1407, and 931-956), the RNAP β N-terminus (residues 1-2), the RNAP α^I^ N-terminus and C-terminal domain (residues 1-5 and 236-329), the RNAP α^II^ N-terminus and C-terminal domain (residues 1-5 and 234-329), the RNAP ω N- and C-termini (residues 1 and 77-91), σ^70^ region 1.1 (residues 1-89), parts of σR2 (residues 166-214, 238-241), σR3 through the σR3-σR4 linker, and parts of σR4 (residues 448-551 and 600-613), density was absent, suggesting high segmental flexibility; these segments were not fitted.

Refinement o the initial model was performed using phenix.real_space_refine under Phenix (12, 29, 30). Each chain was rigid-body refined against the map, and then each chain was subjected to iterative cycles of model building and refinement in Coot (15), followed by real-space refinement with geometry, rotamer, Ramachandran-plot, Cβ, non-crystallographic-symmetry, secondary-structure, and reference-structure (initial model as reference) restraints, followed by global minimization and local rotamer fitting. Secondary-structure annotation was inspected and edited using UCSF Chimera, and geometry was monitored using MolProbity (17). Following model building for each chain, chains were combined and were further refined with geometry, rotamer, Ramachandran-plot, Cβ, non-crystallographic-symmetry, and secondary-structure restraints. The final refined model was validated against over-fitting by randomizing atomic coordinates by 0.3 Å; performing real-space refinement with geometry, rotamer, Ramachandran-plot, Cβ, non-crystallographic-symmetry, secondary-structure, and reference-structure (final refined model as reference) restraints against half-map 1; and performing FSC calculations comparing the resulting model to half-map 1, half-map 2, and the full map (Fig. S3).

Following validation of the final refined model, atom-displacement factors (*B*-factors) were real-space refined against the full map. The final map and atomic model have been deposited in the Electron Microscopy Data Bank (EMDB) and the PDB with accession EMD-21158 and 6VEX, respectively (Figs. 4, S3-S4; Table S2).

#### Cryo-EM structure determination: data collection and data reduction: NusA-containing, engaged Qλ-loading complex

Cryo-EM data for the NusA-containing “engaged” Qλ-loading complex were collected at the Rutgers Cryo-EM and Nanoimaging Facility, using a 200 kV Talos Arctica electron microscope (FEI/ThermoFisher, Inc.) equipped with a K2 Summit direct electron detector (Gatan, Inc.) and a GIF Quantum imaging filter (Gatan, Inc.) with slit width of 20 eV. Data were collected automatically in counting mode using EPU (FEI/ThermoFisher, Inc.), a nominal magnification of 130,000x, a calibrated pixel size of 1.024 Å per pixel, and a dose rate of 6.0 electrons/pixel/s. Movies were recorded at 200 ms/frame for 5 s (25 frames total), resulting in a total radiation dose of 30 electrons/Å^2^. Defocus range was varied between -1.5 µm and -2.5 µm. A total of 1,643 micrographs were recorded from one grid over two days. Micrographs were gain-normalized and defect-corrected.

Data were processed as summarized in Fig. S5A. Dose weighting, motion correction, and CTF estimation were performed as described above for the pre-engaged Qλ-loading complex. Automatic particle picking with Laplacian-of-Gaussian filtering yielded an initial set of 511,891 particles. Particles were extracted into 256x256 pixel boxes, 4x down-scaled, and subjected to rounds of reference-free 2D classification and removal of poorly populated classes, yielding a selected set of 308,619 particles. The selected set was 3D-classified with C1 symmetry using a 60 Å low-pass-filtered map calculated from a cryo-EM structure of an *E. coli* TEC [PDB 6ALF; (22)] as the 3D template, and the best-resolved class, comprising 195,637 particles, was 3D auto-refined and then subjected to 3D classification without alignment, using the auto-refined map as the 3D template. Following 3D classification, the sub-class exhibiting resolved Qλ- QBE density was selected, and the 53,408 particles in the sub-class were re-centered, re-extracted without down-scaling, and subjected to 3D auto-refinement without a reference mask, using the previous auto-refined map as the 3D-template. The refined particles were subjected to 3D classification without alignment, and one subclass (39,378 particles) exhibiting resolved Qλ-QBE density but not NusA density and one subclass (12,931 particles) exhibiting both Qλ-QBE and NusA density were selected. Both subclasses were refined without a mask, and were CTF refined, Bayesian polished (23), further CTF refined, and post-processed. The two subclasses yielded reconstructed maps at 3.2 Å and 3.36 Å overall resolution, respectively, as determined from gold-standard FSC (25, 26), using phase-randomization to account for convolution effects of a solvent mask on FSC between two independently refined half maps, “half-map 1” and “half-map 2” [(27); Fig. S5-S6; Table S2]. The resulting map lacking NusA density (3.2 Å) matched the map of the pre-engaged Qλ-loading complex (Figs. S2-S3; 3.13 Å) and was not used for model building. The resulting map having NusA density (3.36 Å) was used for model building.

The initial atomic model for the NusA-bound engaged Qλ-loading complex was built by manual docking of the cryo-EM structure of the pre-engaged Qλ-loading complex (Figs. 4, S3), and NusA segments from crystal structures of *E. coli* NusA [PDB 5LM7 and 5LM9; (31)] and RNAP α subunit C-terminal domain (αCTD) segments from a crystal structure of the CAP-αCTD-DNA complex [PDB 2O5J; (32)], using UCSF Chimera (24). For the RNAP β’ N and C-termini and SI3 (residues 1-14, 1376-1407, and 931-956), the RNAP β N-terminus (residues 1-3), the RNAP α^I^ N-terminus and C-terminal domain (residues 1-5 and 236-329), the RNAP α^II^ N-terminus, linker, and part of the C-terminal domain (residues 1-5, 234-249 and 316-329), the RNAP ω N- and C-termini (residues 1 and 77-91), σ^70^ region 1.1 (residues 1-89), parts of σR2 (residues 166-214 and 238-241), σR3 through the σR3-σR4 linker and part of σR4 (residues 448-551 and 600-613), and the NusA KH2, AR1, and AR2 domains (residues 341-495), density was absent, suggesting high segmental flexibility; these segments were not fitted. Refinement and validation were performed as described above for the pre-engaged Qλ-loading complex. The final map and atomic model were deposited in the EMDB and the PDB with accession codes EMD-21159 and 6VEY, respectively (Figs. 5, S5-S6; Table S2).

## SUPPLEMENTAL FIGURE LEGENDS

**Fig. S1.**
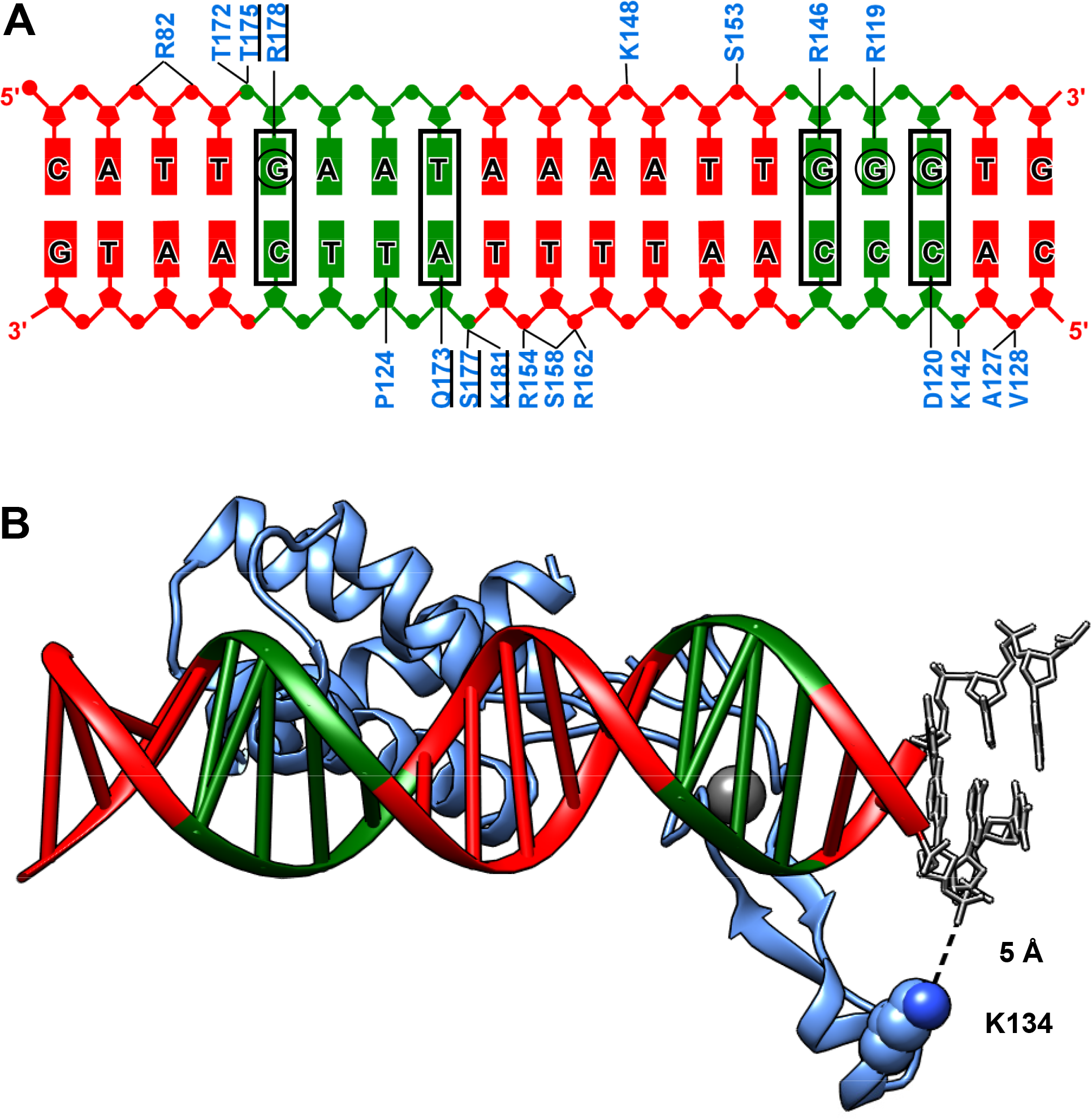
Structure of Qλ-QBE complex: Qλ-QBE interactions. **(A)** Summary of Qλ-QBE interactions. Qλ residues at which alanine substitution results in decreased Qλ-QBE interaction are underlined (1), base pairs at which substitutions result in decreased Qλ-QBE interaction are boxed (1, 33), and guanine bases that exhibit methylation-protection or methylation-enhancement in DNA-footprinting experiments, indicative of protein-DNA interaction with the DNA major groove, are circled (34). **(B)** Location of Qλ K134 relative to DNA. Modeled extension of QBE DNA fragment, gray. Other colors and view orientation as in Fig. 3A, left.

**Fig. S2.**
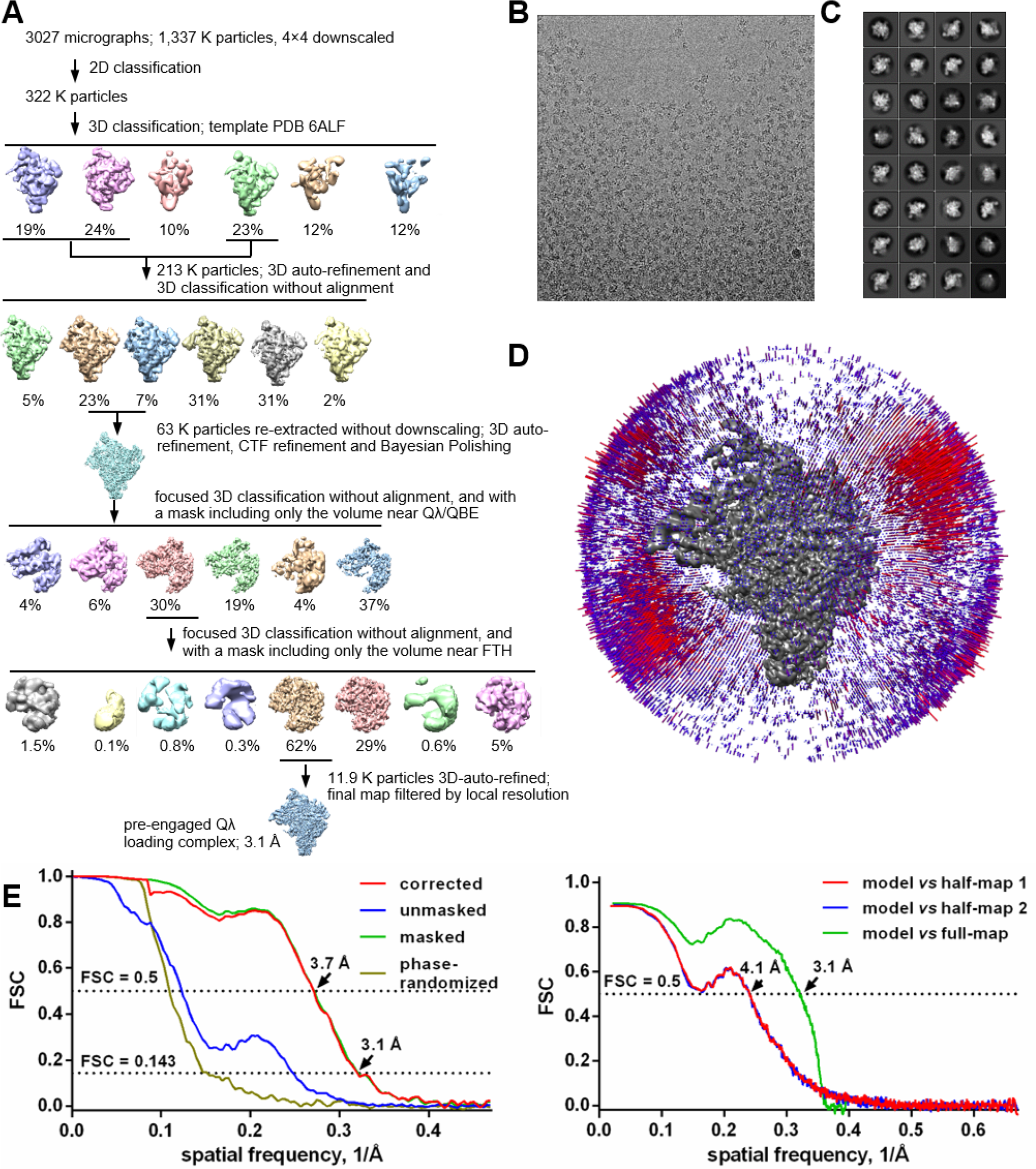
Structure of pre-engaged Qλ-loading complex: structure determination. **(A)** Data processing scheme. **(B)** Representative electron micrograph. **(C)** 2D-class averages. **(D)** Euler-angle distribution. **(E)** Fourier-shell-correlation (FSC) plots.

**Fig. S3.**
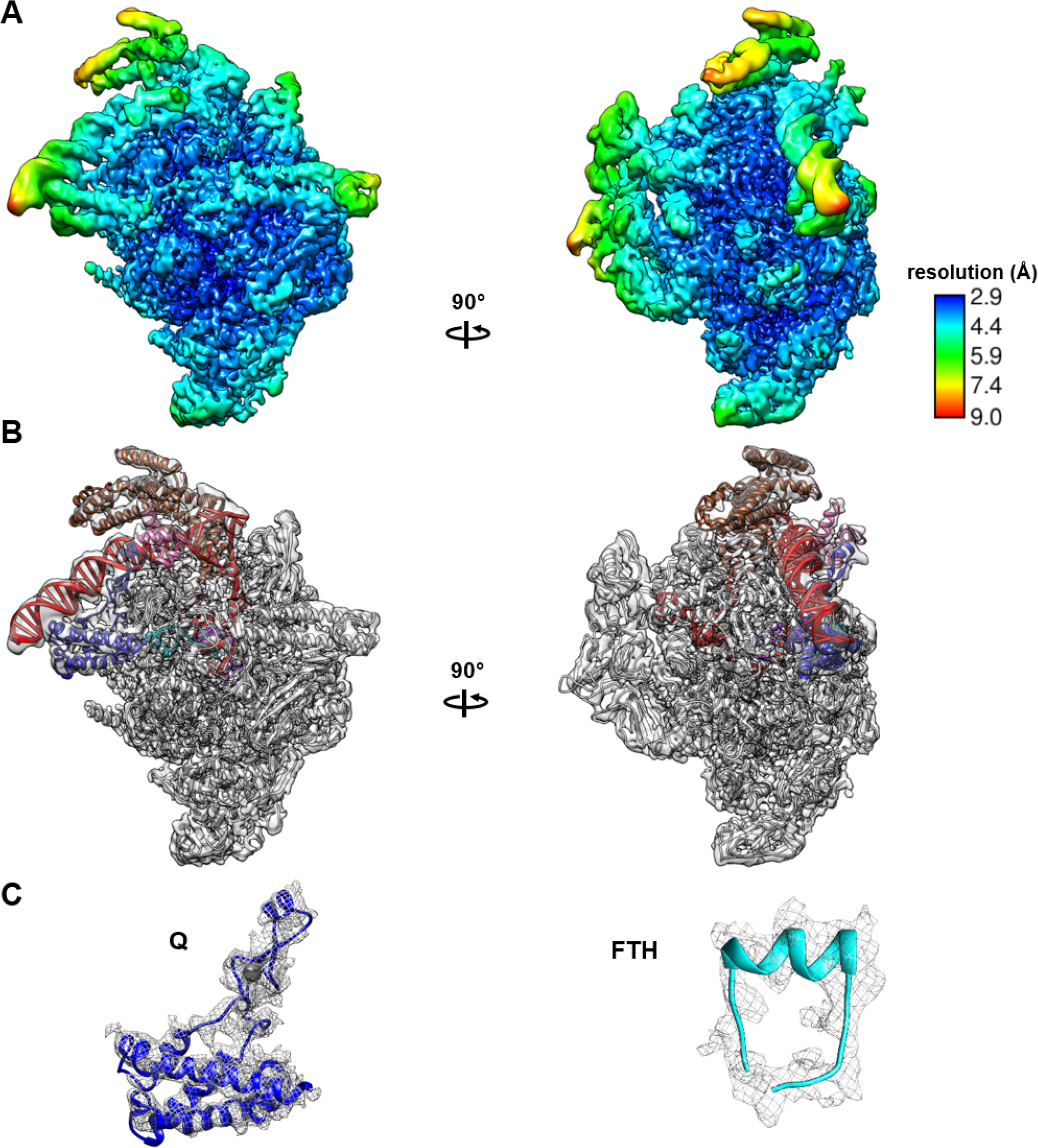
Structure of pre-engaged Qλ-loading complex: maps and fit. **(A)** EM density map colored by local resolution. View orientations as in Fig. 4B. **(B)** EM-density map and fit. View orientations and colors as in Fig. 4B. **(C)** Density maps for Qλ (left) and RNAP FTH (right).

**Fig. S4.**
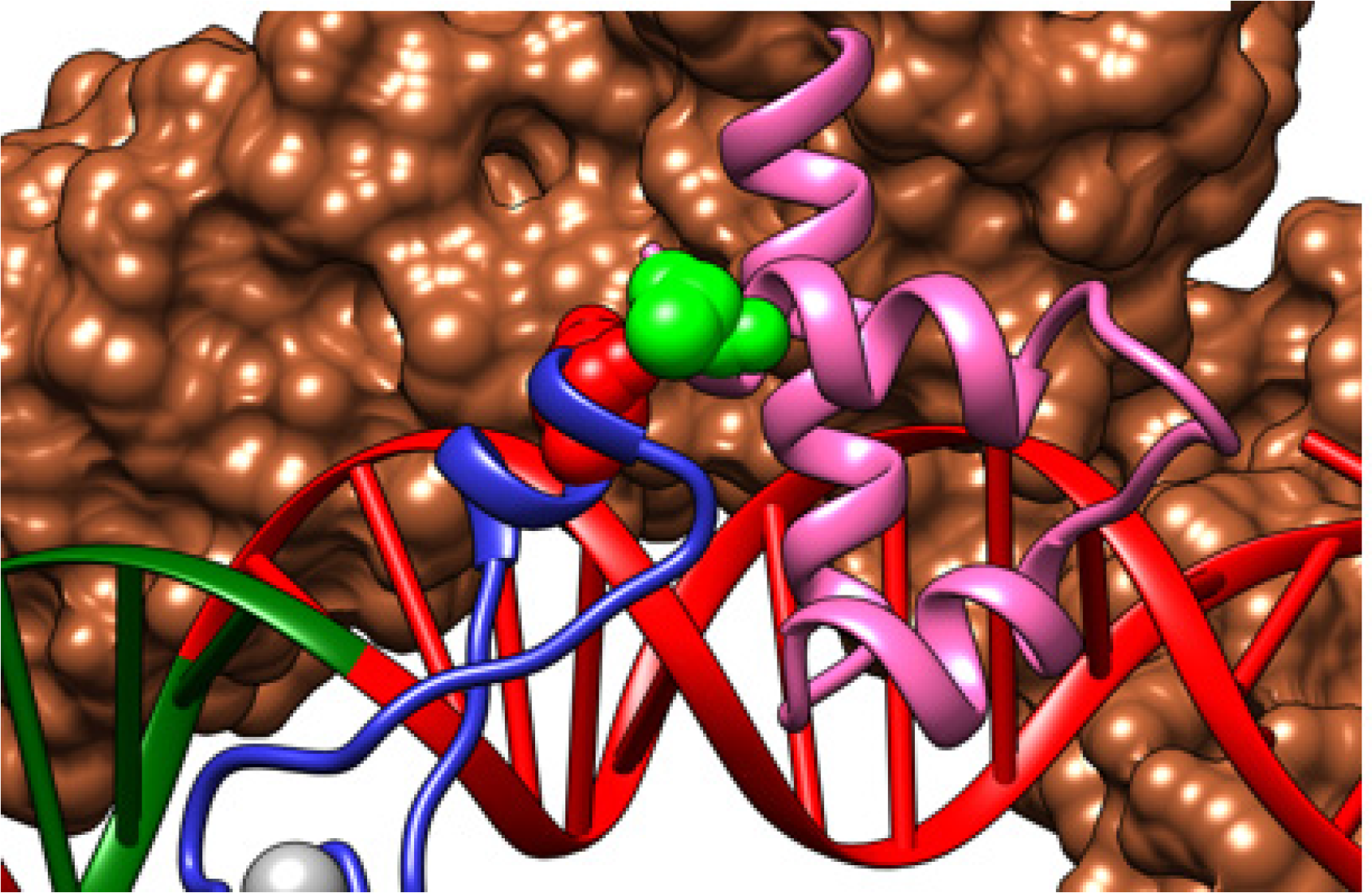
Structure of “pre-engaged” Qλ-loading complex: repositioning of σR4. Interactions of Qλ arm (blue) with σR4 (pink) repositioned to DNA segment immediately upstream of σR2 (brown surface) on SDPE. Site of substitution of Qλ that results in defect in Qλ-σR4 interaction is shown in red [E134; (13)], and site of substitution of σR4 that results in defect in Qλ-σR4 interaction is shown in green [A553; (35)].

**Fig. S5.**
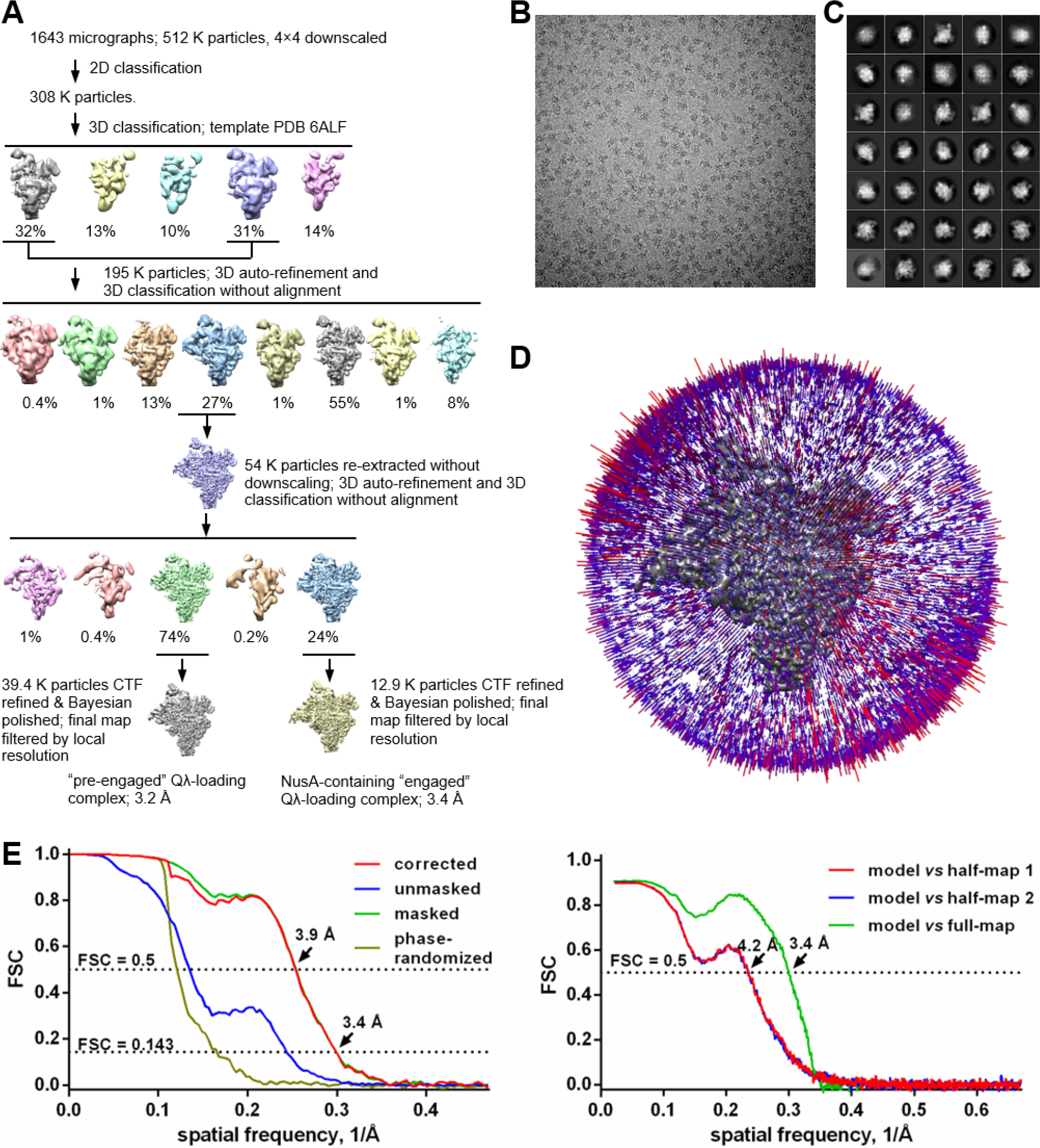
Structure of NusA-containing, engaged Qλ-loading complex: structure determination. **(A)** Data processing scheme. **(B)** Representative electron micrograph. **(C)** 2D-class averages. **(D)** Euler-angle distribution. **(E)** Fourier-shell-correlation (FSC) plots.

**Fig. S6.**
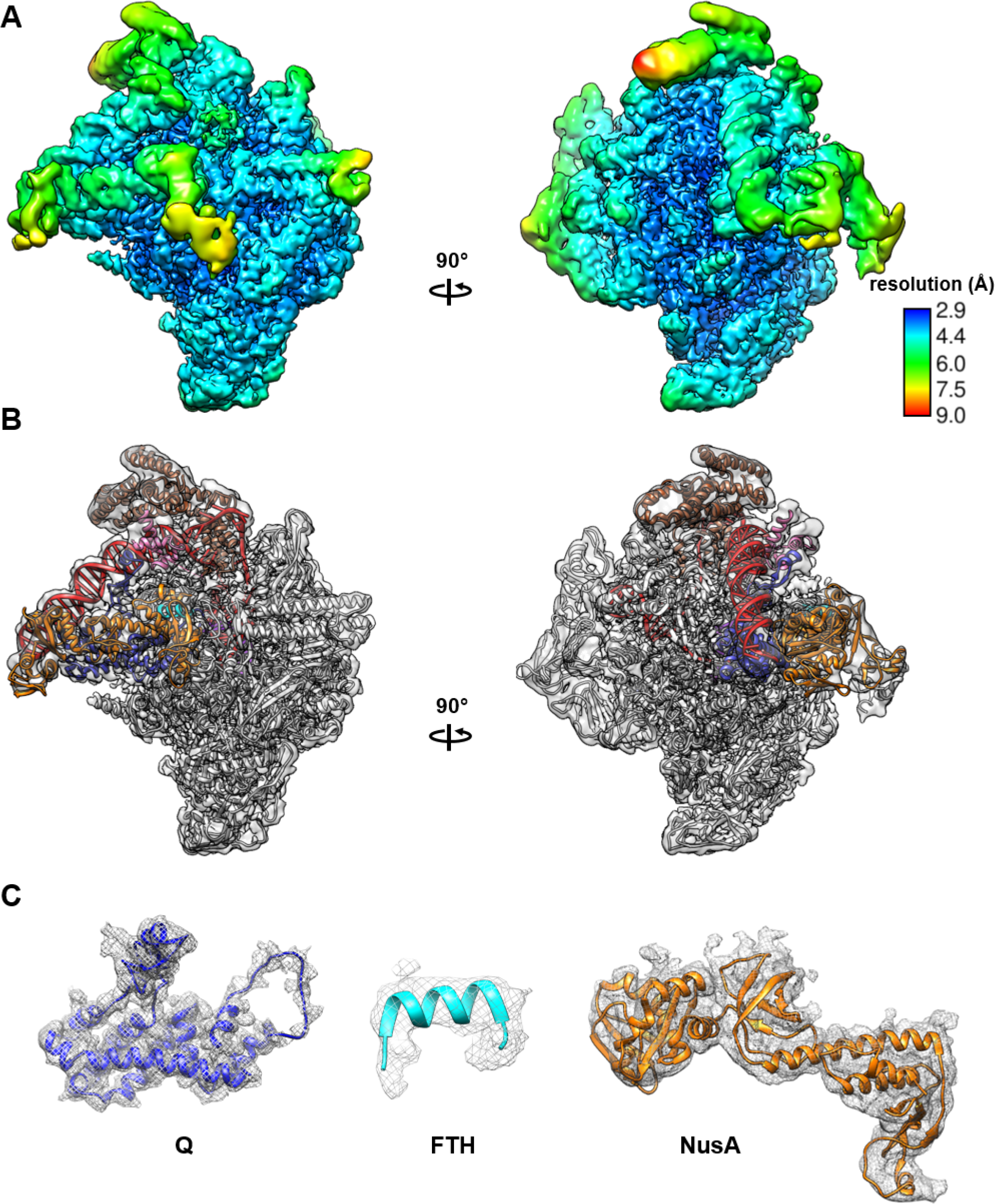
Structure of NusA-containing, engaged Qλ-loading complex: maps and fit. **(A)** EM density map colored by local resolution. View orientations as in Figs. 4B and 5A. **(B)** EM-density map and fit. View orientations and colors as in Figs. 4B and 5A. **(C)** Density maps for Qλ (left), RNAP FTH (center), and NusA (right).

**Fig. S7.**
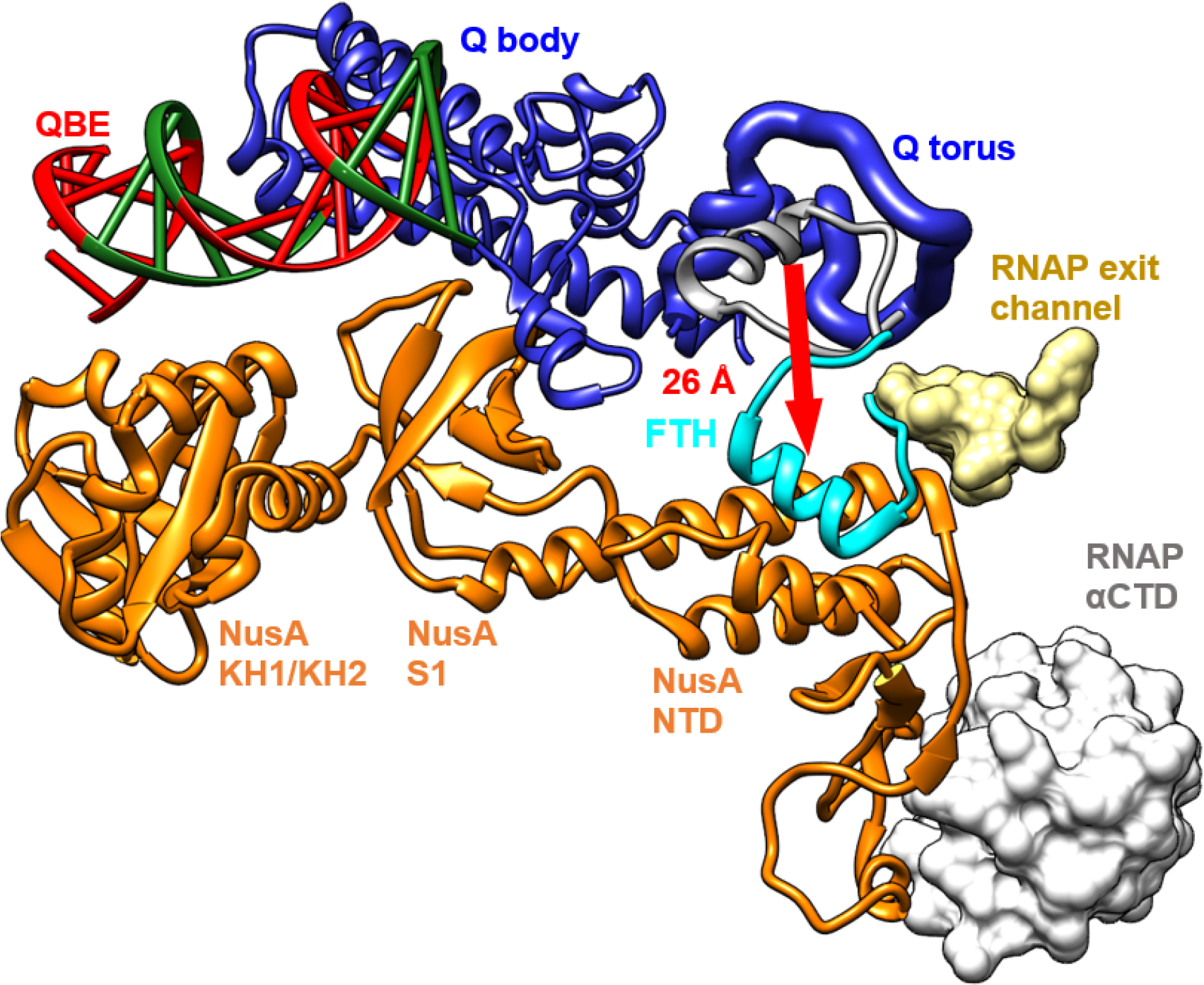
Structure of NusA-containing, engaged Qλ-loading complex: NusA-Qλ and NusA-RNAP interactions. RNAP FTH in NusA-containing, engaged Qλ-loading complex, cyan; RNAP FTH in pre-engaged Qλ-loading complex, medium gray; RNAP β subunit residues at rim of RNAP RNA-exit channel, light yellow; and RNAP α CTD, light gray. Other colors as in Fig. 5A. Red arrow shows the 26 Å re-positioning of RNAP FTH to vacate space to be occupied by the Qλ torus.

**Fig. S8.**
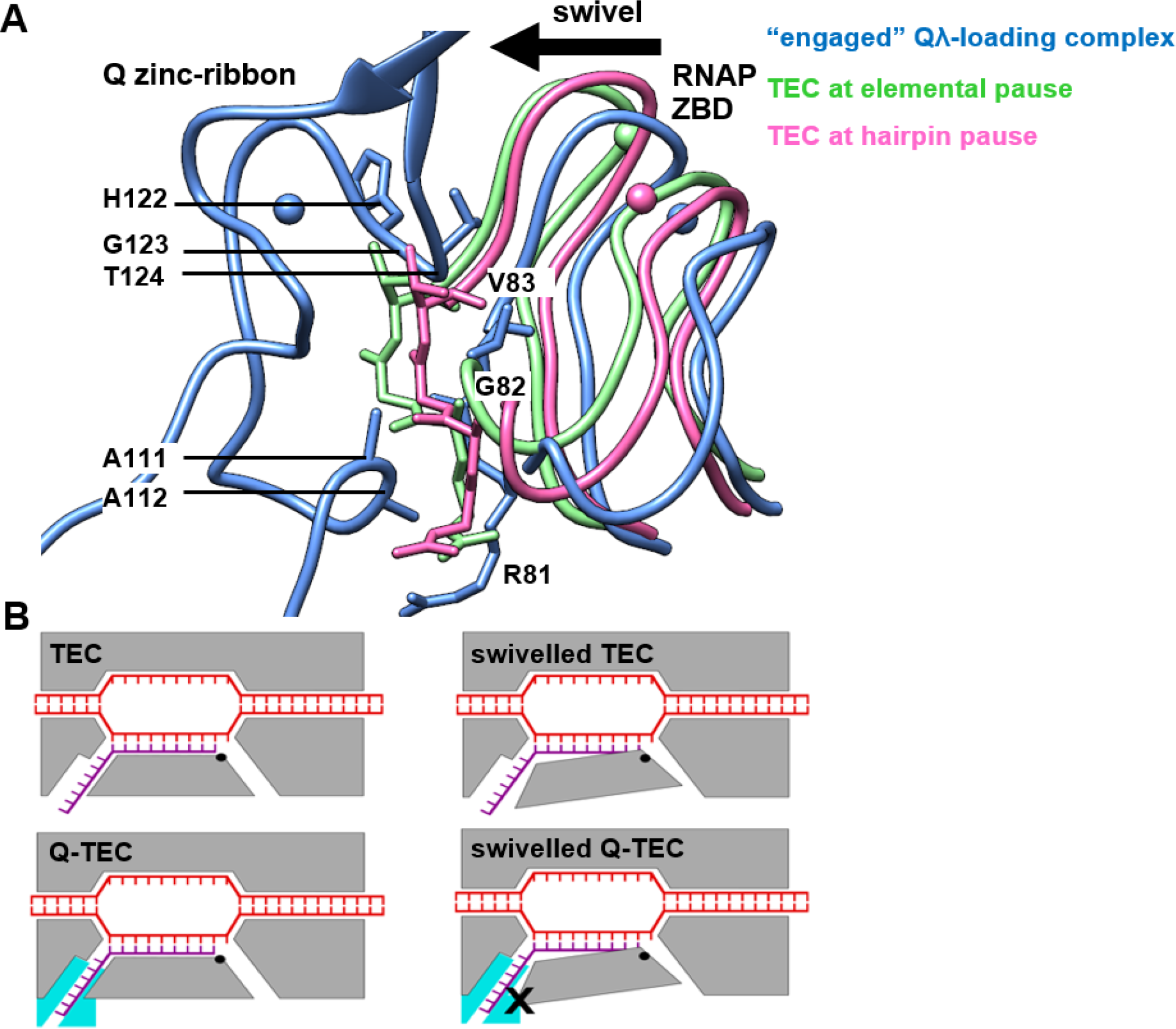
Suppression of RNAP swivelling by Qλ. **(A)** Interactions of Qλ zinc ribbon with RNAP ZBD that preclude swivelling. Three structures are superimposed: NusA-containing, engaged Qλ-loading complex (blue), TEC with swivelling of ∼3° associated with hairpin-dependent pausing [PDB 6ASX; (36); see also (37); pink], and TEC with swivelling of ∼1.5° associated with elemental pausing [PDB 6BJS; (36) green]. Movement of RNAP ZBD upon RNAP swivelling is indicated with arrow. RNAP swivelling is expected to decrease distance and create steric clash between the Qλ zinc ribbon and the RNAP ZBD. **(B)** Schematic comparison of TECs in absence of Qλ (upper row) and TECs in presence of Qλ (lower row). Steric clash between Qλ zinc ribbon and RNAP ZBD in the swivelled state, X. Colors as in Fig. 4B and 5A.

## SUPPLEMENTAL MOVIES

**Movie S1.** Comparison of Qλ conformation in free Qλ (type-I crystal structure) to Qλ conformation in Qλ-QBE. Colors and view orientation as in Fig. 3B, left.

**Movie S2.** Comparison of Qλ conformation in Qλ-QBE (gray) to Qλ conformation in pre-engaged Qλ- loading complex (blue). View orientation as in Fig. 4E.

**Movie S3.** Comparison of Qλ conformation in pre-engaged Qλ-loading complex (gray) to Qλ conformation in NusA-containing, engaged Qλ-loading complex (blue). View orientation as in Fig. 5D.

**Table S1.**
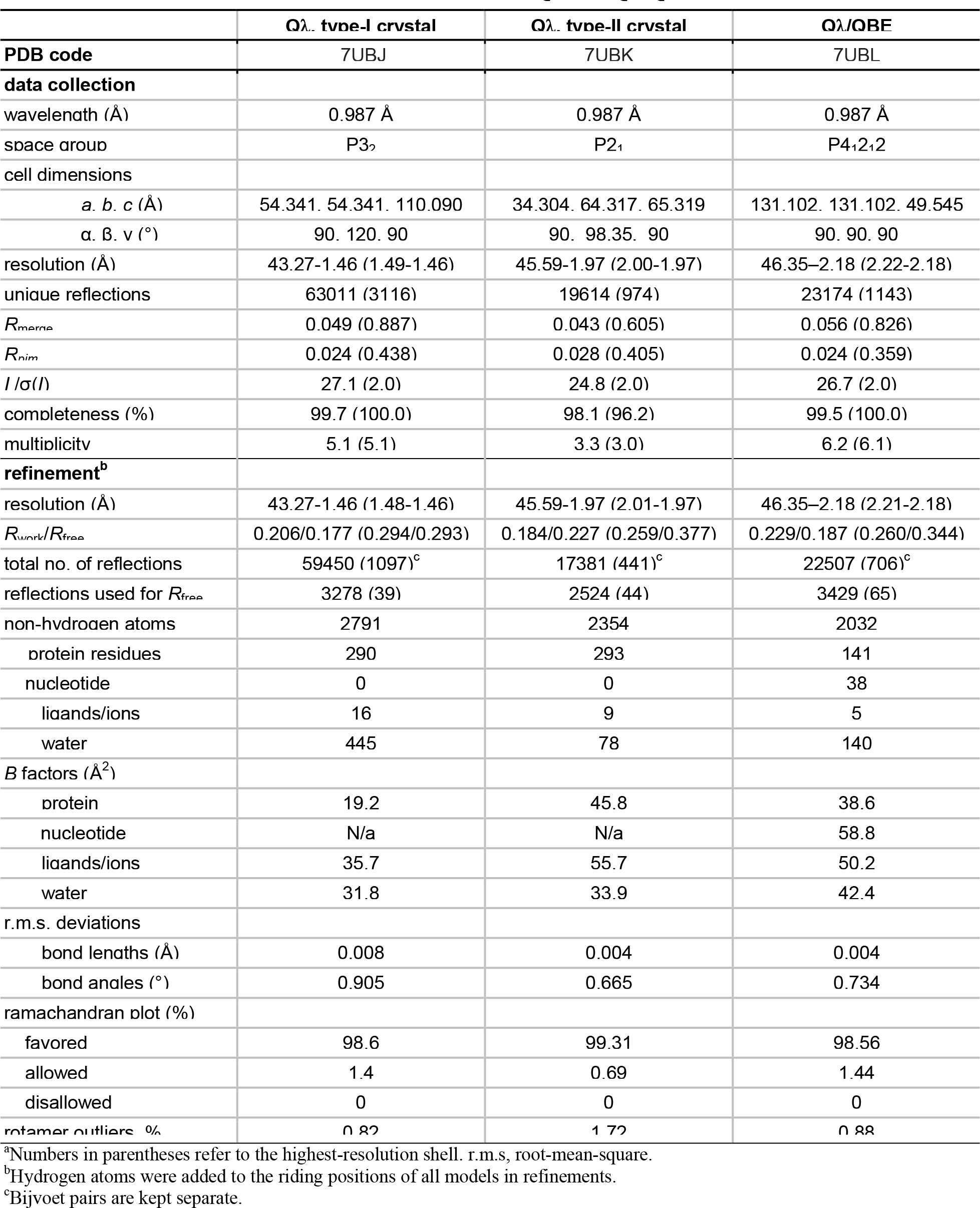
Data collection and refinement statistics for Qλ and Qλ-QBE.^a^.

**Table S2.** Data-collection, refinement, and validation statistics for cryo-EM structures of pre-engaged Qλ-loading complex and NusA-containing, engaged Qλ-loading complex.

| | pre-engaged Qλ-loading complex | NusA-containing, engaged Qλ-loading complex |
| --- | --- | --- |
| **PDB code** | 7UBM | 7UBN |
| **EMDB code** | EMD-26438 | EMD-26439 |
| **data collection and processing** |  |  |
| microscope | FEI Titan Krios | FEI Talos Arctica |
| camera | Gatan K2-Summit | Gatan K2-Summit |
| energy Filter | N/a | 20 ev |
| magnification (nominal) | 47,608x (22,500x) | 130,000x |
| voltage (kV) | 300 | 200 |
| electron exposure (e-/Å<sup>2</sup>) | 56.2 | 30 |
| defocus range (μm) | -0.8 to -1.5 | -1.5 to -2.5 |
| pixel size (Å) | 0.533 | 1.024 |
| symmetry imposed | C1 | C1 |
| micrographs (no.) | 3,027 | 1,643 |
| initial particle images (no.) | 1,337,270 | 511,891 |
| final particle images (no.) | 11,913 | 12,931 |
| map resolution (Å) | 3.13 | 3.36 |
| FSC threshold | 0.143 | 0.143 |
| map resolution range (Å) | 999-3.13 | 999-3.36 |
| **refinement** |  |  |
| initial model used (PDB code) | 6P18, 4YLN, Qλ/QBE crystal structure | Pre-engaged Qλ-loading complex, 5LM7, 5LM9, 1LB2 |
| model resolution (Å) | 3.13 | 3.36 |
| FSC threshold | 0.143 | 0.143 |
| model resolution range (Å) | 46.9-3.13 | 49.5-3.36 |
| map sharpening *B* factor (Å<sup>2</sup>) | -25 | -22 |
| model composition |  |  |
| non-hydrogen atoms | 31,584 | 35,070 |
| protein residues | 3716 | 4165 |
| nucleotide | 116 | 116 |
| ligands | 4 | 4 |
| *B* factors (Å<sup>2</sup>) |  |  |
| protein | 91.7 | 99.9 |
| Nucleotide | 147.0 | 133.8 |
| ligand | 100.8 | 110.7 |
| r.m.s. deviations |  |  |
| bond lengths (Å) | 0.005 | 0.006 |
| bond angles (°) | 0.800 | 0.849 |
| validation |  |  |
| molProbity score | 2.10 | 2.31 |
| clashscore | 6.15 | 7.36 |
| poor rotamers (%) | 4.11 | 5.7 |
| Ramachandran plot |  |  |
| favored (%) | 95.75 | 94.80 |
| allowed (%) | 4.25 | 5.20 |
| disallowed (%) | 0.00 | 0.00 |

## Notes

### Competing Interest Statement

The authors have declared no competing interest.

## REFERENCES

1. J. W. Roberts, Transcription termination and late control in phage lambda. Proc Natl Acad Sci U S A 72, 3300–3304 (1975).

2. E. J. Grayhack, J. W. Roberts, The phage lambda Q gene product: activity of a transcription antiterminator *in vitro*. Cell 30, 637–648 (1982).

3. E. J. Grayhack, X. J. Yang, L. F. Lau, J. W. Roberts, Phage lambda gene Q antiterminator recognizes RNA polymerase near the promoter and accelerates it through a pause site. Cell 42, 259–269 (1985).

4. W. S. Yarnell, J. W. Roberts, The phage lambda gene Q transcription antiterminator binds DNA in the late gene promoter as it modifies RNA polymerase. Cell 69, 1181–1189 (1992).

5. B. Z. Ring, J. W. Roberts, Function of a nontranscribed DNA strand site in transcription elongation. Cell 78, 317–324 (1994).

6. B. Z. Ring, W. S. Yarnell, J. W. Roberts, Function of *E. coli* RNA polymerase sigma factor sigma 70 in promoter-proximal pausing. Cell 86, 485–493 (1996).

7. W. S. Yarnell, J. W. Roberts, Mechanism of intrinsic transcription termination and antitermination. Science 284, 611–615 (1999).

8. J. W. Roberts, W. Yarnell, E. Bartlett, J. Guo, M. Marr, D. C. Ko, H. Sun, C. W. Roberts, Antitermination by bacteriophage lambda Q protein. Cold Spring Harb Symp Quant Biol 63, 319–325 (1998).

9. R. A. Weisberg, M. E. Gottesman, Processive antitermination. J Bacteriol 181, 359–367 (1999).

10. J. W. Roberts, S. Shankar, J. J. Filter, RNA polymerase elongation factors. Annu Rev Microbiol 62, 211–233 (2008).

11. P. Deighan, A. Hochschild, The bacteriophage λQ anti-terminator protein regulates late gene expression as a stable component of the transcription elongation complex. Mol Microbiol 63, 911–920 (2007).

12. Z. Yin, J. T. Kaelber, R. H. Ebright, Structural basis of Q-dependent antitermination. Proc Natl Acad Sci U S A 116, 18384–18390 (2019).

13. H. C. Guo, M. Kainz, J. W. Roberts, Characterization of the late-gene regulatory region of phage 21. J Bacteriol 173, 1554–1560 (1991).

14. W. Yarnell, Interaction of the antitermination factor Q with complexes of RNA polymerase and DNA. Ph.D. dissertation, Cornell University (1993).

15. X. J. Yang, J. A. Goliger, J. W. Roberts, Specificity and mechanism of antitermination by Q proteins of bacteriophages lambda and 82. J Mol Biol 210, 453–460 (1989).

16. J. Shi, X. Gao, T. Tian, Z. Yu, B. Gao, A. Wen, L. You, S. Chang, X. Zhang, Y. Zhang, Y. Feng, Structural basis of Q-dependent transcription antitermination. Nat Commun 10, 2925 (2019).

17. S. M. Vorobiev, Y. Gensler, H. Vahedian-Movahed, J. Seetharaman, M. Su, J. Y. Huang, R. Xiao, G. Kornhaber, G. T. Montelione, L. Tong, R. H. Ebright, B. E. Nickels, Structure of the DNA-binding and RNA-polymerase-binding region of transcription antitermination factor λQ. Structure 22, 488–495 (2014).

18. J. Guo, J. W. Roberts, DNA binding regions of Q proteins of phages λ and phi80. J Bacteriol 186, 3599–3608 (2004).

19. C. Andreini, I. Bertini, G. Cavallaro, Minimal functional sites allow a classification of zinc sites in proteins. PLoS One 6, e26325 (2011).

20. E. Bartlett, Characterization of the functional interaction between the bacteriophage λ Q antiterminator and late gene promoter DNA. Ph.D. dissertation, Cornell University (1998).

21. B. E. Nickels, S. J. Garrity, V. Mekler, L. Minakhin, K. Severinov, R. H. Ebright, A. Hochschild, The interaction between σ^70^ and the β-flap of *Escherichia coli* RNA polymerase inhibits extension of nascent RNA during early elongation. Proc Natl Acad Sci U S A 102, 4488–4493 (2005).

22. B. E. Nickels, C. W. Roberts, J. W. Roberts, A. Hochschild, RNA-mediated destabilization of the σ^70^ region 4/β-flap interaction facilitates engagement of RNA polymerase by the Q antiterminator. Mol Cell 24, 457–468 (2006).

23. P. Deighan, C. M. Diez, M. Leibman, A. Hochschild, B. E. Nickels, The bacteriophage λ Q antiterminator protein contacts the β-flap domain of RNA polymerase. Proc Natl Acad Sci U S A 105, 15305–15310 (2008).

24. C. Pukhrambam, V. Molodtsov, M. Kooshkbaghi, A. Tareen, H. Vu, K. S. Skalenko, M. Su, Z. Yin, J. T. Winkelman, J. B. Kinney, R. H. Ebright, B. E. Nickels, Structural and mechanistic basis of σ dependent transcriptional pausing. *Proc Natl Acad Sci U S A* (under review; https://www.biorxiv.org/content/10.1101/2022.01.24.477500v1) (2022).

25. D. G. Vassylyev, S. Sekine, O. Laptenko, J. Lee, M. N. Vassylyeva, S. Borukhov, S. Yokoyama, Crystal structure of a bacterial RNA polymerase holoenzyme at 2.6 A resolution. Nature 417, 712–719 (2002).

26. K. S. Murakami, S. Masuda, E. A. Campbell, O. Muzzin, S. A. Darst, Structural basis of transcription initiation: an RNA polymerase holoenzyme-DNA complex. Science 296, 1285–1290 (2002).

27. V. Mekler, E. Kortkhonjia, J. Mukhopadhyay, J. Knight, A. Revyakin, A. N. Kapanidis, W. Niu, Y. W. Ebright, R. Levy, R. H. Ebright, Structural organization of bacterial RNA polymerase holoenzyme and the RNA polymerase-promoter open complex. Cell 108, 599–614 (2002).

28. K. Kuznedelov, L. Minakhin, A. Niedziela-Majka, S. L. Dove, D. Rogulja, B. E. Nickels, A. Hochschild, T. Heyduk, K. Severinov, A role for interaction of the RNA polymerase flap domain with the σ subunit in promoter recognition. Science 295, 855–857 (2002).

29. D. A. Siegele, J. C. Hu, W. A. Walter, C. A. Gross, Altered promoter recognition by mutant forms of the σ^70^ subunit of *Escherichia coli* RNA polymerase. J Mol Biol 206, 591–603 (1989).

30. Y. Zuo, T. A. Steitz, Crystal structures of the *E. coli* transcription initiation complexes with a complete bubble. Mol Cell 58, 534–540 (2015).

31. B. Bae, A. Feklistov, A. Lass-Napiorkowska, R. Landick, S. A. Darst, Structure of a bacterial RNA polymerase holoenzyme open promoter complex. eLife 4 (2015).

32. M. T. Marr, S. A. Datwyler, C. F. Meares, J. W. Roberts, Restructuring of an RNA polymerase holoenzyme elongation complex by lambdoid phage Q proteins. Proc Natl Acad Sci U S A 98, 8972–8978 (2001).

33. B. E. Nickels, C. W. Roberts, H. Sun, J. W. Roberts, A. Hochschild, The σ^70^ subunit of RNA polymerase is contacted by the λQ antiterminator during early elongation. Mol Cell 10, 611-622 (2002).

34. P. G. Devi, E. A. Campbell, S. A. Darst, B. E. Nickels, Utilization of variably spaced promoter- like elements by the bacterial RNA polymerase holoenzyme during early elongation. Mol Microbiol 75, 607–622 (2010).

35. J. Guo, Analysis of the functional domains and characterization of the DNA-binding domain of phage λ Q protein. Ph.D. dissertation, Cornell University (1999).

36. R. S. Washburn, M. E. Gottesman, Regulation of transcription elongation and termination. Biomolecules 5, 1063–1078 (2015).

37. G. A. Belogurov, I. Artsimovitch, Regulation of transcript elongation. Annu Rev Microbiol 69, 49–69 (2015).

38. J. Y. Kang, T. V. Mishanina, M. J. Bellecourt, R. A. Mooney, S. A. Darst, R. Landick, RNA polymerase accommodates a pause RNA hairpin by global conformational rearrangements that prolong pausing. Mol Cell 69, 802–815 e805 (2018).

39. X. Guo, A. G. Myasnikov, J. Chen, C. Crucifix, G. Papai, M. Takacs, P. Schultz, A. Weixlbaumer, Structural basis for NusA stabilized transcriptional pausing. Mol Cell 69, 816–827 e814 (2018).

40. F. Krupp, N. Said, Y. H. Huang, B. Loll, J. Burger, T. Mielke, C. M. T. Spahn, M. C. Wahl, Structural basis for the action of an all-purpose transcription anti-termination factor. Mol Cell 74, 143–157 e145 (2019).

41. C. Wang, V. Molodtsov, E. Firlar, J. T. Kaelber, G. Blaha, M. Su, R. H. Ebright, Structural basis of transcription-translation coupling. Science 369, 1359–1365 (2020).

42. N. Said, T. Hilal, N. D. Sunday, A. Khatri, J. Burger, T. Mielke, G. A. Belogurov, B. Loll, R. Sen, I. Artsimovitch, M. C. Wahl, Steps toward translocation-independent RNA polymerase inactivation by terminator ATPase rho. Science 371 (2021).

43. Z. Hao, V. Epshtein, K. H. Kim, S. Proshkin, V. Svetlov, V. Kamarthapu, B. Bharati, A. Mironov, T. Walz, E. Nudler, Pre-termination transcription complex: structure and function. Mol Cell 81, 281–292 e288 (2021).

44. C. Zhu, X. Guo, P. Dumas, M. Takacs, M. Abdelkareem, A. Vanden Broeck, C. Saint-André, G. Papai, C. Crucifix, J. Ortiz, A. Weixlbaumer, Transcription factors modulate RNA polymerase conformational equilibrium. Nat Commun 13, 1546 (2022).

45. K. S. Murakami, S. Masuda, S. A. Darst, Structural basis of transcription initiation: RNA polymerase holoenzyme at 4 Å resolution. Science 296, 1280–1284 (2002).

46. A. Ray-Soni, M. J. Bellecourt, R. Landick, Mechanisms of bacterial transcription termination. Annu Rev Biochem 85, 319–347 (2016).

47. J. W. Roberts, Mechanisms of bacterial transcription termination. J Mol Biol 431, 4030–4039 (2019).

48. J. Y. Kang, T. V. Mishanina, R. Landick, S. A. Darst, Mechanisms of transcriptional pausing in bacteria. J Mol Biol 10.1016/j.jmb.2019.07.017 (2019).

49. G. A. Belogurov, I. Artsimovitch, The mechanisms of substrate selection, catalysis, and translocation by the elongating RNA polymerase. J Mol Biol 431, 3975–4006 (2019).

50. R. Landick, Transcriptional pausing as a mediator of bacterial gene regulation. Annu Rev Microbiol 75, 291–314 (2021).

51. C. D. Wells, P. Deighan, M. Brigham, A. Hochschild, Nascent RNA length dictates opposing effects of NusA on antitermination. Nucleic Acids Res 44, 5378–5389 (2016).

52. S. Shankar, A. Hatoum, J. W. Roberts, A transcription antiterminator constructs a NusA- dependent shield to the emerging transcript. Mol Cell 27, 914–927 (2007).

## REFERENCES

1. J. Guo, J. W. Roberts, DNA binding regions of Q proteins of phages lambda and phi80. J Bacteriol 186, 3599–3608 (2004).

2. Z. Yin, J. T. Kaelber, R. H. Ebright, Structural basis of Q-dependent antitermination. Proc Natl Acad Sci U S A 116, 18384–18390 (2019).

3. J. Guo, Analysis of the functional domains and characterization of the DNA-binding domain of phage lambda Q protein. Ph.D. dissertation, Cornell University (1999).

4. M. T. Marr, J. W. Roberts, Function of transcription cleavage factors GreA and GreB at a regulatory pause site. Mol Cell 6, 1275–1285 (2000).

5. V. Svetlov, I. Artsimovitch, Purification of bacterial RNA polymerase: tools and protocols. Methods Mol Biol 1276, 13–29 (2015).

6. B. E. Nickels, S. J. Garrity, V. Mekler, L. Minakhin, K. Severinov, R. H. Ebright, A. Hochschild, The interaction between σ70 and the β-flap of Escherichia coli RNA polymerase inhibits extension of nascent RNA during early elongation. Proceedings of the National Academy of Sciences 102, 4488–4493 (2005).

7. B. E. Nickels, C. W. Roberts, J. W. Roberts, A. Hochschild, RNA-mediated destabilization of the sigma(70) region 4/beta flap interaction facilitates engagement of RNA polymerase by the Q antiterminator. Mol Cell 24, 457–468 (2006).

8. P. Deighan, C. M. Diez, M. Leibman, A. Hochschild, B. E. Nickels, The bacteriophage lambda Q antiterminator protein contacts the beta-flap domain of RNA polymerase. Proc Natl Acad Sci U S A 105, 15305–15310 (2008).

9. G. Panaghie, S. E. Aiyar, K. L. Bobb, R. S. Hayward, P. L. de Haseth, Aromatic amino acids in region 2.3 of Escherichia coli sigma 70 participate collectively in the formation of an RNA polymerase-promoter open complex. J Mol Biol 299, 1217–1230 (2000).

10. P. G. Devi, E. A. Campbell, S. A. Darst, B. E. Nickels, Utilization of variably spaced promoter- like elements by the bacterial RNA polymerase holoenzyme during early elongation. Mol Microbiol 75, 607–622 (2010).

11. Z. Otwinowski, W. Minor, Processing of X-ray diffraction data collected in oscillation mode. Methods Enzymol 276, 307–326 (1997).

12. P. D. Adams, P. V. Afonine, G. Bunkoczi, V. B. Chen, I. W. Davis, N. Echols, J. J. Headd, L. W. Hung, G. J. Kapral, R. W. Grosse-Kunstleve, A. J. McCoy, N. W. Moriarty, R. Oeffner, R. J. Read, D. C. Richardson, J. S. Richardson, T. C. Terwilliger, P. H. Zwart, PHENIX: a comprehensive Python-based system for macromolecular structure solution. Acta Crystallogr D Biol Crystallogr 66, 213–221 (2010).

13. S. M. Vorobiev, Y. Gensler, H. Vahedian-Movahed, J. Seetharaman, M. Su, J. Y. Huang, R. Xiao, G. Kornhaber, G. T. Montelione, L. Tong, R. H. Ebright, B. E. Nickels, Structure of the DNA-binding and RNA-polymerase-binding region of transcription antitermination factor lambdaQ. Structure 22, 488–495 (2014).

14. B. C. Wang, Resolution of phase ambiguity in macromolecular crystallography. Methods Enzymol 115, 90–112 (1985).

15. P. Emsley, K. Cowtan, Coot: model-building tools for molecular graphics. Acta Crystallogr D Biol Crystallogr 60, 2126–2132 (2004).

16. J. Painter, E. A. Merritt, Optimal description of a protein structure in terms of multiple groups undergoing TLS motion. Acta Crystallogr D Biol Crystallogr 62, 439–450 (2006).

17. V. B. Chen, W. B. Arendall, 3rd, J. J. Headd, D. A. Keedy, R. M. Immormino, G. J. Kapral, L. W. Murray, J. S. Richardson, D. C. Richardson, MolProbity: all-atom structure validation for macromolecular crystallography. Acta Crystallogr D Biol Crystallogr 66, 12–21 (2010).

18. B. Carragher, N. Kisseberth, D. Kriegman, R. A. Milligan, C. S. Potter, J. Pulokas, A. Reilein, Leginon: an automated system for acquisition of images from vitreous ice specimens. J Struct Biol 132, 33–45 (2000).

19. S. Q. Zheng, E. Palovcak, J. P. Armache, K. A. Verba, Y. Cheng, D. A. Agard, MotionCor2: anisotropic correction of beam-induced motion for improved cryo-electron microscopy. Nat Methods 14, 331–332 (2017).

20. K. Zhang, Gctf: Real-time CTF determination and correction. J Struct Biol 193, 1–12 (2016).

21. J. Zivanov, T. Nakane, B. O. Forsberg, D. Kimanius, W. J. Hagen, E. Lindahl, S. H. Scheres, New tools for automated high-resolution cryo-EM structure determination in RELION-3. Elife 7 (2018).

22. J. Y. Kang, P. D. Olinares, J. Chen, E. A. Campbell, A. Mustaev, B. T. Chait, M. E. Gottesman, S. A. Darst, Structural basis of transcription arrest by coliphage HK022 Nun in an Escherichia coli RNA polymerase elongation complex. Elife 6 (2017).

23. J. Zivanov, T. Nakane, S. H. W. Scheres, A Bayesian approach to beam-induced motion correction in cryo-EM single-particle analysis. IUCrJ 6, 5–17 (2019).

24. E. F. Pettersen, T. D. Goddard, C. C. Huang, G. S. Couch, D. M. Greenblatt, E. C. Meng, T. E. Ferrin, UCSF Chimera--a visualization system for exploratory research and analysis. J Comput Chem 25, 1605–1612 (2004).

25. S. Chen, G. McMullan, A. R. Faruqi, G. N. Murshudov, J. M. Short, S. H. Scheres, R. Henderson, High-resolution noise substitution to measure overfitting and validate resolution in 3D structure determination by single particle electron cryomicroscopy. Ultramicroscopy 135, 24–35 (2013).

26. R. Henderson, A. Sali, M. L. Baker, B. Carragher, B. Devkota, K. H. Downing, E. H. Egelman, Z. Feng, J. Frank, N. Grigorieff, W. Jiang, S. J. Ludtke, O. Medalia, P. A. Penczek, P. B. Rosenthal, M. G. Rossmann, M. F. Schmid, G. F. Schroder, A. C. Steven, D. L. Stokes, J. D. Westbrook, W. Wriggers, H. Yang, J. Young, H. M. Berman, W. Chiu, G. J. Kleywegt, C. L. Lawson, Outcome of the first electron microscopy validation task force meeting. Structure 20, 205–214 (2012).

27. S. H. Scheres, S. Chen, Prevention of overfitting in cryo-EM structure determination. Nat Methods 9, 853–854 (2012).

28. Y. Zuo, T. A. Steitz, Crystal structures of the E. coli transcription initiation complexes with a complete bubble. Molecular cell 58, 534–540 (2015).

29. R. P. D. Adams, R. W. Grosse-Kunstleve, L. W. Hung, T. R. Ioerger, A. J. McCoy, N. W. Moriarty, R. J. Read, J. C. Sacchettini, N. K. Sauter, T. C. Terwilliger, PHENIX: building new software for automated crystallographic structure determination. Acta Crystallogr D Biol Crystallogr 58, 1948–1954 (2002).

30. P. V. Afonine, B. K. Poon, R. J. Read, O. V. Sobolev, T. C. Terwilliger, A. Urzhumtsev, P. D. Adams, Real-space refinement in PHENIX for cryo-EM and crystallography. Acta Crystallogr D Struct Biol 74, 531–544 (2018).

31. N. Said, F. Krupp, E. Anedchenko, K. F. Santos, O. Dybkov, Y. H. Huang, C. T. Lee, B. Loll, E. Behrmann, J. Burger, T. Mielke, J. Loerke, H. Urlaub, C. M. T. Spahn, G. Weber, M. C. Wahl, Structural basis for lambdaN-dependent processive transcription antitermination. Nat Microbiol 2, 17062 (2017).

32. B. Benoff, H. Yang, C. L. Lawson, G. Parkinson, J. Liu, E. Blatter, Y. W. Ebright, H. M. Berman, R. H. Ebright, Structural basis of transcription activation: the CAP-alpha CTD-DNA complex. Science 297, 1562–1566 (2002).

33. W. S. Yarnell, J. W. Roberts, The phage lambda gene Q transcription antiterminator binds DNA in the late gene promoter as it modifies RNA polymerase. Cell 69, 1181–1189 (1992).

34. E. Bartlett, Characterization of the functional interaction between the bacteriophage lambda Q antiterminator and late gene promoter DNA. (1998).

35. B. E. Nickels, C. W. Roberts, H. Sun, J. W. Roberts, A. Hochschild, The sigma(70) subunit of RNA polymerase is contacted by the (lambda)Q antiterminator during early elongation. Mol Cell 10, 611–622 (2002).

36. S. J. Y. Kang, R. A. Mooney, Y. Nedialkov, J. Saba, T. V. Mishanina, I. Artsimovitch, R. Landick, A. Darst, Structural basis for transcript elongation control by NusG family universal regulators. Cell 173, 1650–1662 e1614 (2018).

37. X. Guo, A. G. Myasnikov, J. Chen, C. Crucifix, G. Papai, M. Takacs, P. Schultz, A. Weixlbaumer, Structural basis for NusA stabilized transcriptional pausing. Mol Cell 69, 816–827 e814 (2018).

